# Neuronal and non-neuronal functions of the synaptic cell adhesion molecule, neurexin, in *Nematostella vectensis*

**DOI:** 10.1101/2024.04.09.588470

**Authors:** Christine Guzman, Kurato Mohri, Yuko Tsuchiya, Kentaro Tomii, Hiroshi Watanabe

## Abstract

The transition from diffusion-mediated cell-cell communication to faster and more targeted synaptic signaling in animal nervous systems has long been of interest to evolutionary biologists. Although genome sequencing of early-diverging metazoans has revealed the broad distribution of synapse-related genes among them, synaptic structures have been observed only in Cnidaria, the sister group to Bilateria. How synaptic machinery evolved remains largely unknown. In this study, we investigated the function of neurexins (Nrxns), a core family of presynaptic cell adhesion molecules with critical roles in bilaterian chemical synapses, using the cnidarian model, *Nematostella vectensis*. Neural Nrxns, named delta-Nrxns, are expressed mainly in neuronal cell clusters that exhibit both peptidergic and classical neurotransmitter signaling. Knockdown of Nrxnδ genes reduced spontaneous peristalsis of *N. vectensis* polyps. Interestingly, gene knockdown and pharmacological studies suggested that Nrxnδ is involved in glutamate- and glycine-mediated signaling rather than peptidergic signaling. Knockdown of the epithelial Nrxn in *N. vectensis* revealed a major role in cell adhesion, particularly between ectodermal and endodermal epithelia. Overall, this study provides molecular, functional, and cellular insights into the ancestral, non-neural function of Nrxns, as well as key information for understanding how and why this family of cell adhesion molecules was recruited to synaptic machinery.

## INTRODUCTION

The nervous system takes center stage among key innovations in animal evolution. Its appearance drastically shaped how animals perceive signals from the environment, integrate and store information, and mount appropriate responses. Chemical synapses are one of the major modes of network-dependent signaling in the nervous system. In recent years, comparative genomic analyses across various metazoan species have revealed the conservation and deep evolutionary origins of synapse constituent genes. Many proteins that have synaptic functions in Bilateria are found in non-bilaterian genomes (Putnam et al. 2007; Ryan et al. 2013; Moroz et al. 2014), including neuron-less animals (Srivastava et al. 2008; Srivastava et al. 2010; Musser et al. 2021), and in unicellular close relatives of metazoans (Sakarya et al. 2007; Suga et al. 2013; Burkhardt et al. 2014). It remains a mystery, however, how synapse-dependent neural signaling emerged early during animal evolution and how synaptic protein machinery was later refined.

During the early evolution of neural systems, diffusion-based volume transmission by peptides may have been the primary mode of cell–cell communication (Senatore et al. 2017; Takahashi 2020; Jekely 2021; Sachkova et al. 2021; Hayakawa et al. 2022). Later, faster and more targeted synaptic signaling was needed, as animal bodies and behaviors increased in complexity (Keijzer et al. 2017). One key factor that sets chemical synaptic signaling apart from diffusible signaling is the establishment of precise cell–cell contacts between the presynaptic neurons and the postsynaptic cells. These contacts are facilitated and stabilized by Synaptic Adhesion Molecules (SAMs). Several SAMs are also involved in assembly of intracellular protein complexes, effectively contributing to maintenance, specificity, and plasticity of synapses (Dalva et al. 2007; Rudenko 2017; Sudhof 2018). Disruption of SAMs cause synaptic dysfunction, leading to various neurodegenerative and neurodevelopmental diseases (Gorlewicz et al. 2018; Leshchyns’ka et al. 2016).

Among all SAMs, neurexins (Nrxns) are major presynaptic hub molecules at chemical synapses in Bilateria (Gomez et al. 2021; Sudhof 2017, 2021). Depending on which postsynaptic ligand is bound, Nrxns may promote excitatory or inhibitory synapse development (Graf et al. 2004). Furthermore, Nrxns are involved in Ca^2+^-triggered neurotransmitter release (Missler et al. 2003) and proper organization of synaptic molecules (Hata et al. 1996; Li et al. 2007; Zhang et al. 2010; Trotter et al. 2019). Alterations of Nrxn gene sequence or expression have been linked to neuropsychiatric and neurodegenerative disorders (Craig et al. 2007; Sudhof 2017). Previous studies reported that Nrxns are expressed mainly in neurons and astrocytes (Gokce et al. 2013; Uchigashima et al. 2019). However, several phylum-wide gene surveys demonstrated that Nrxns predated neuron and synapse appearance and that they are conserved in all metazoan lineages (Ganot et al. 2015; Moroz et al. 2015), suggesting that Nrxns served functions unrelated to synapses in early metazoan ancestors. Still, neither pre-neural functions of Nrxns nor why they became major components of neural synapses during metazoan evolution are well known.

In this paper, we first compared structures of Nrxn proteins and their features required for synaptic function, mainly targeting early divergent lineages of metazoans. Interestingly, bilaterians and cnidarians are the only lineages in which Nrxns are abundantly expressed in the nervous system. In ctenophores, the earliest branching lineage with a nervous system, Nrxns are expressed in a wide range of cell types, including epithelial cells, as they are in neuron-less poriferans and placozoans. Then, we examined functions of the two types of Nrxns — neuronal and non-neuronal Nrxns — in the sea anemone, *Nematostella vectensis*, a cnidarian that is the closest outgroup to the Bilateria. We show that neuronal Nrxns are involved in motor control of *N. vectensis* polyps and that this function is independent of neuropeptide-mediated neurotransmission. Pharmacological testing suggested instead the involvement of small-molecule neurotransmitters, glutamate and glycine. Furthermore, we observed that the widely expressed and more ancestral non-neuronal Nrxn is involved in proper heterophilic cell-cell adhesion. Altogether, these results enhance our understanding of putative synaptically mediated neurotransmission in non-bilaterians and their non-neuronal ancestral functions.

## RESULTS

### Identification and characterization of non-bilaterian Nrxns

At synapses in bilaterians, Nrxn exerts its functions by binding to various proteins through its intracellular and extracellular domains (Figure 1a). To explore possible roles of Nrxns during early nervous system evolution, we first surveyed the presence of Nrxn homologs and their domain organization in metazoans and their closest unicellular relatives. In contrast to a previous phylogenetic study of Nrxn superfamily members (Ganot et al. 2015), we performed a detailed comparison focused on the organization of Laminin/Neurexin/Sex hormone-binding globulin (LNS) repeated domains. Despite low sequence identity (20%), LNS domains show high structural similarity as revealed by the crystal structure of Nrxns (Miller et al. 2011). Therefore, their structural information is useful for obtaining ideal alignments between phylogenetically distant bilaterian and prebilaterian Nrxns sequences. We first performed a structural prediction based on *Bos taurus* alpha-Nrxn1 structures (PDB IDs: 3QCW and 3POY) and found a conserved LNS domain organization in prebilaterian and bilaterian Nrxns (Figure 1b). Our molecular phylogenetic tree using alignments based on structural information confirmed the presence of Nrxn genes in all metazoan species surveyed, with the exception of some poriferan species (Figure 1c, Supplementary Figure 1,2). Bilaterian Nrxns grouped in a single clade with bilaterian Casprs (contactin-associated proteins), also members of the Nrxn superfamily. However, no non-bilaterian Nrxn-like genes contained the N-terminal discoidin domain typical of Caspr proteins (Supplementary Figure 2). Hence, it seems that Caspr genes first appeared in the bilaterian lineage.

**Figure 1:**
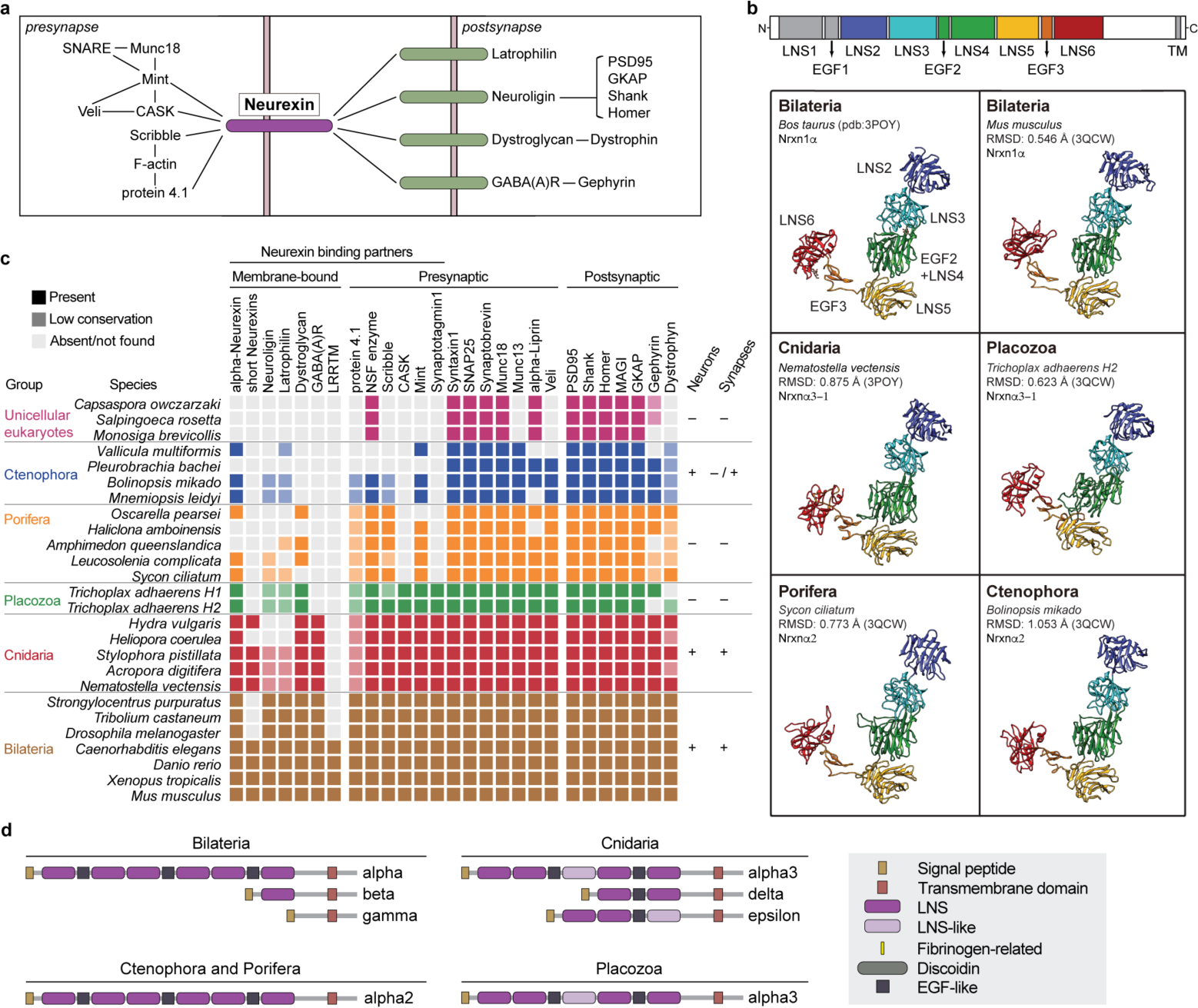
Phylogenetic distribution and domain organization of neurexins. **a** Overview of synaptic protein complexes mediated by Nrxns in Bilateria. **b** 3D structure models of bilaterian and non-bilaterian Nrxns. Domain organization of bovine α-Neurexin 1 (PDB ID:3POY) used as a template for homology modelling (upper panel). The crystal structure of bovine NRXN1A includes seven contiguous domains of the extracellular region (LNS2 to LNS6 domains and intervening EGF2 and EGF3 domains) (lower panels, top left). Predicted 3D structures of Nrxns of *Mus musculus* (Bilateria), *Nematostella vectensis* (Cnidaria), *Bolinopsis mikado* (Ctenophora), *Trichoplax adhaerens H2* (Placozoa), and *Sycon ciliatum* (Porifera) are shown. The root-mean-square deviation (RSMD) values in Angstroms between the template and modelled structures were also calculated. **c** Distribution of Neurexin (Nrxn), its binding partners, and other associated synaptic proteins in metazoan lineages and their closest unicellular relatives. **d** Domain organizations of the Nrxn proteins. Bilaterian α-Nrxn (full-length) consists of an N-terminal signal peptide, followed by three repeats of LNS(A)-EGF-LNS(B), a single transmembrane domain, and a short cytoplasmic tail. Shorter splice variants (β-Nrxn (vertebrates only) and γ-Nrxn) of bilaterian Nrxns are also shown. Non-bilaterian Nrxns are classified into four types: α2-, α3-, δ-, and ε-Nrxns.

A survey of known binding partners of Nrxns demonstrated that many postsynaptic interacting factors, e.g., Dystroglycan, appeared after the last common ancestor of Placozoa/Cnidaria/Bilateria (Figure 1c). Meanwhile, among the major Nrxn partner candidates, only Latrophilin and Neuroligin have homologs in the Ctenophora and Porifera. The paucity of Nrxn apparatus components in early-branching metazoans contrasts with the full deployment of other membrane signaling proteins, such as Semaphorin/Plexin and Ephrin/EphrinR (Supplementary Figure 3).

There are three Nrxn genotypes in vertebrates, named α, β, and γ, but only α and β types contain LNS domains. Therefore, γ-Nrxn was excluded from our analysis. All investigated members of Porifera (other than demosponges *Amphimedon queenslandica* and *Haliclona amboinensis*) and Ctenophora have a single Nrxn homolog, “α2-Nrxn”, belonging to the α-type. Poriferan and ctenophoran α2-Nrxns possess six LNS domains and three epidermal growth factor (EGF) domains, similar to the full-length α-Nrxn of bilaterians. Cnidarians and placozoans also harbor an α-like Nrxn, named here as “α3-Nrxn”, which contains only four or five LNS domains (Figure 1d, Supplementary Figure 1). In addition to the α3-Nrxns, cnidarians also possess other Nrxn types. Firstly, a new type, named here as “δ-Nrxn”, is a short Nrxn sequence that is observed only in cnidarians. This type is a reminiscent of the short β and γ isoforms found in bilaterians. Secondly, a type named “ε-Nrxn” is supposedly a derivative of the α3-Nrxns. However, some regions, in particular at the C-terminus, are rather divergent and cannot be annotated with any protein domains. Unlike the β-Nrxns in bilaterians, which are produced by alternative splicing from the same gene as α-Nrxn, the cnidarian α-, ε-, and δ-Nrxns are encoded by separate genes. Thus, we refer to them as different types rather than isoforms. Interestingly, based on our phylogenetic tree, the cnidarian δ-Nrxns are sister group to bilaterian Nrxns and Casprs (Supplementary Figure 2). On the other hand, all non-bilaterian α3-Nrxns formed a monophyletic group together with cnidarian-specific ε-Nrxns. These findings suggest that the α-type of Nrxn is the most ancestral, and that the ε and α types were derived from a common ancestral gene. It is still unknown why short-type Nrxns expanded uniquely in cnidarians, but the diversity of the domain organization and expression signatures of these Nrxns may be caused by increased Nrxn functional requirements (discussed below). It is remarkable that the two most conserved LNS domains (LNS#2 and LNS#6) across all species are the only domains in Nrxns known to participate in binding with postsynaptic targets (Supplementary Figure 1). The preservation of the core domain, in contrast to the increased Nrxn gene repertoire in prebilaterians, suggests evolutionary conservation in the mode of action of Nrxns, including partner binding.

To gain insight into the function of Nrxn proteins in early metazoans, we next examined the expression profile of Nrxn genes in known cell types (Sebe-Pedros, Chomsky, et al. 2018; Sebe-Pedros, Saudemont, et al. 2018). In *N vectensis*, δ-Nrxns (NvNrxnδ1,-2,-3, and -4) and ε-Nrxns (NvNrxnε1, -2, and -3) are highly enriched in neurons, while the two homologs of α3-Nrxns (NvNrxnα3-1 and -2) are broadly expressed in different cell types including epithelial cells (Figure 2a, Supplementary Figure 4). Likewise, the α3-Nrxns of *Trichoplax adhaerens* (Placozoa) and α2-Nrxn of *Mnemiopsis leidyi* (Ctenophora) also show broad expression patterns (Supplementary Figure 5). From this point onward, δ- and ε-type Nrxns with nervous system-specific expression will be referred to as neural Nrxns (nNrxns), and widely expressed Nrxns with a more conserved domain organization will be referred to as classical Nrxns (cNrxns). Notably, the PDZ-binding motif at the C-terminus of bilaterian Nrxns crucial for intracellular signaling (Biederer et al. 2000; Hata et al. 1996) is absent in cNrxns of Porifera and Ctenophora (Supplementary Figure 1).

**Figure 2:**
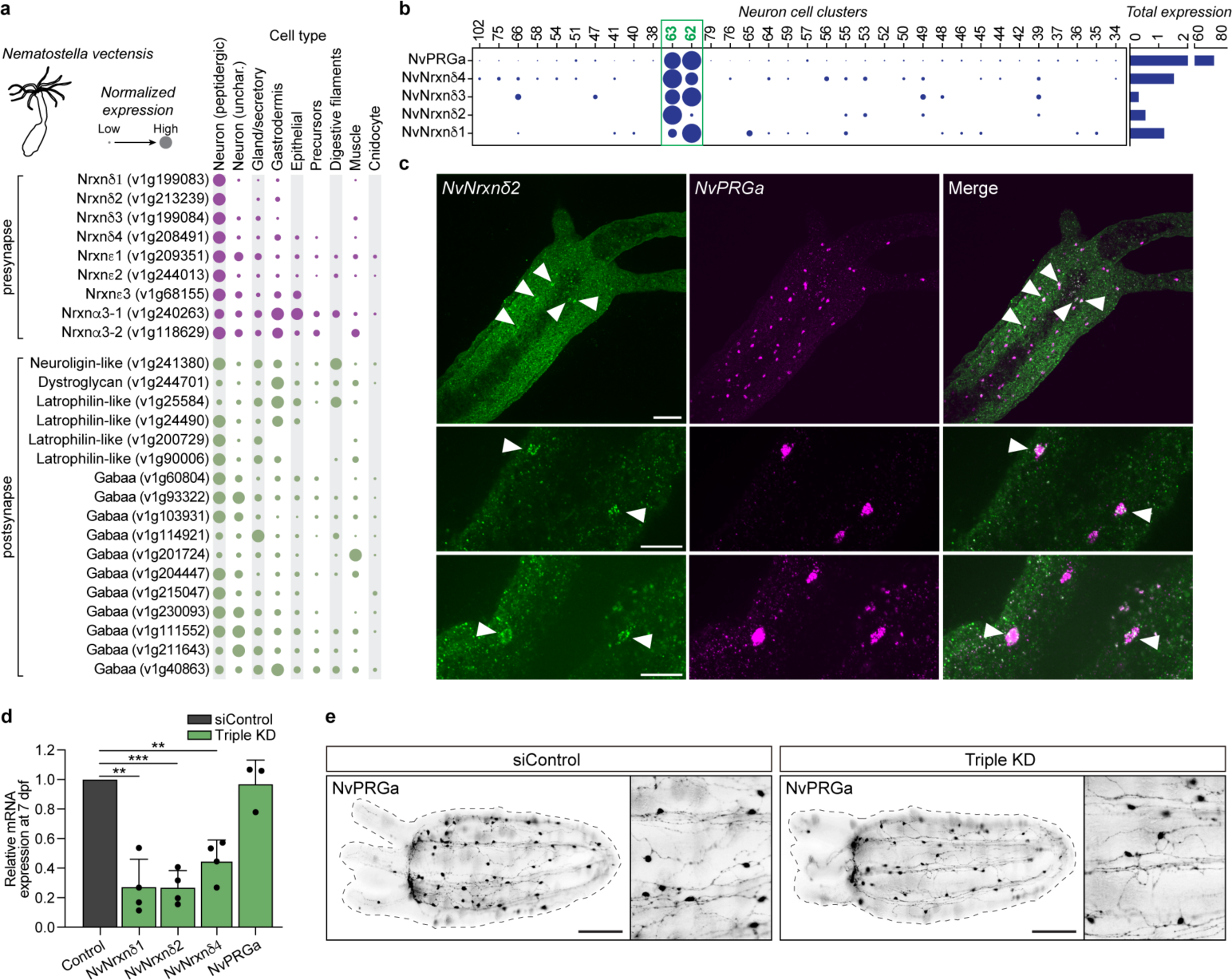
Specific expression of δ-Nrxns in PRGamide-positive neurons. **a** Abundance of Nrxn homologs across cell type categories of adult *Nematostella vectensis*. **b** Abundance of δ-Nrxns across neuronal cell clusters within neuron categories (peptidergic and uncharacterized). Cell clusters 62 and 63 (green box) express both δ-Nrxns and neuropeptide NvPRGamide. **c** Representative images of hybridization chain reaction staining of NvNrxnδ2 and NvPRGamide mRNAs in primary polyps (7 dpf). Arrowheads denote the neurons expressing both NvNrxnδ2 and NvPRGamide genes. **d** qPCR analysis of δ-Nrxns and NvPRGa expression at 7 dpf after triple knockdown of δ-Nrxn genes (NvNrxnδ1, -2, -4). Shown is the mean ± SD; ** and *** denote the statistical significance *p*<0.01 and *p*<0.001 in two-sided unpaired t-test of at least three independent experiments, respectively. **e** Representative images of NvPRGamide-positive neurons visualized by immunostaining of 7-dpf polyps treansfected with control siRNA (left) or siRNAs for NvNrxnδ1 (#2 + #3), NvNrxnδ2 (#3), and NvNrxnδ4 (#1 + #2) (right). In **a** and **b**, dots correspond to normalized UMI count abundance scaled per gene. Dots scale from smallest to largest, corresponding to lowest and highest expression, respectively. Scale bars in **c** top, **c** middle and bottom and **e**, 50, 20 and 100 µm, respectively.

### A population of PRGamide-expressing neurons is nNrxn-positive

We then analyzed whether non-bilaterian Nrxns and their putative membrane-bound partners are expressed in neurons using single-cell expression data. As with Nrxns, many putative partners are expressed in neurons of adult *N. vectensis* (Figure 2a, Supplementary Figure 4). In *T. adhaerens*, expression is observed in epithelial and lipophil cells, as well as in some peptide-expressing cells (Supplementary Figure 5). In *M. leidyi*, Latrophilin-like and Neuroligin-like genes, the only Nrxn partner candidates, are mainly localized in non-neuronal cells, consistently with the broad expression profile of the ctenophore cNrxn. Among the nNrxns of *N. vectensis*, δ genes (NvNrxnδ1,-2,-3, and -4) are enriched in neuronal clusters C62 and C63, suggesting a specific requirement for this type of Nrxns in functions of this subpopulation of neurons (Figure 2b).

Remarkably, the C62/C63 neuronal cluster also exclusively expresses the PRGamide neuropeptide. We then focused on NvNrxnd1,-2, and -4, since NvNrxnδ3 has an incomplete C-terminus. Expressions of NvNrxnδ1,-2, and -4 increased during the period when neural networks develop after the planula stage (Supplementary Figure 6) and are expressed in neurons in polyp stage (Figure 2a,b, Supplementary Figure 4). Since we were not successful in producing specific antibodies to detect NvNrxnδ1,-2, and -4 proteins, we then performed mRNA detection by *in situ* hybridization chain reaction (HCR). Although expression of nNrxn mRNAs was weak, clear expression of NvNrxnδ2 could be detected in at least some PRGamide-expressing neurons (Figure 2c).

### Neuron development is not affected by nNrxn knockdown

In bilaterians, Nrxns are expressed early in development long before synapse formation begins and they are required for neurite development in *Drosophila melanogaster* (Constance et al. 2018). To verify Nrxn function in cnidarians, we first tested the effects of Nrxn knockdown on the developmental process of δ-Nrxn-positive neurons. We performed small interfering RNA (siRNA)-mediated gene knockdown by delivering siRNAs targeting each δ-Nrxn mRNA into *N. vectensis* embryos (Masuda-Ozawa et al. 2022b). Loss of any of the three δ-Nrxns (NvNrxnδ1,-2, and -4) was non-lethal, and caused no visible defects in polyp development (Supplementary Figure 7). The effect of knockdown of δ-Nrxns on development of the neural network was further evaluated by immunostaining for PRGamide in the primary polyps. Single-gene knockdown showed no obvious changes in development, projection patterning, and distribution of the δ-Nrxn^+^ neurons (Supplementary Figure 8). The expression level of the PRGamide gene was also unaffected. To test the possibility that loss of a single δ-Nrxn could be compensated by functional redundancy by other δ-Nrxn proteins, we performed triple gene knockdown targeting all three δ-Nrxns. However, even in embryos in which all three nNrxns were knocked down, no significant defects in neuron development and assembly were observed, nor was deregulation of PRGamide gene expression (Figure 2d,e). These results suggest that cnidarian δ-Nrxns do not contribute to development of the PRGamide^+^ neuronal network.

### nNrxn knockdown impairs locomotor activity

In bilateral animals, such as mice and *Drosophila*, where Nrxn functions as a synaptic member, loss of Nrxns negatively influences locomotor activity (Grayton et al. 2013; Li et al. 2007). Similarly, we carried out behavioral assays to assess whether the reduction of nNrxns has the same effect in *N. vectensis*. Among the several behaviors of *N. vectensis* polyps, we observed and quantified peristalsis, a rhythmic radial contraction of the body column of the polyp that starts from the region just below the pharynx and propagates to the foot. This peristaltic movement, which is actively observed even in petri dishes, is a major spontaneous behavior of burrowing sea anemone polyps. Knockdown of a single nNrxn gene significantly decreased the number of peristaltic waves (Figure 3a). Similarly, triple-knockdown polyps displayed fewer waves compared to control animals, but unexpectedly, it did not elicit further wave reduction compared to single-knockdown animals (Figure 3a-c). A previous study showed that reduced peristaltic waves in juvenile *N. vectensis* is correlated to a reduced number of neurons (Havrilak et al. 2021), indicating that peristalsis is under the control of neural activities. Our results suggest that the function of δ-Nrxns expressed in specific PRGamide^+^ neuronal subpopulation play an essential role in the neuronal activities responsible for controlling peristalsis.

**Figure 3:**
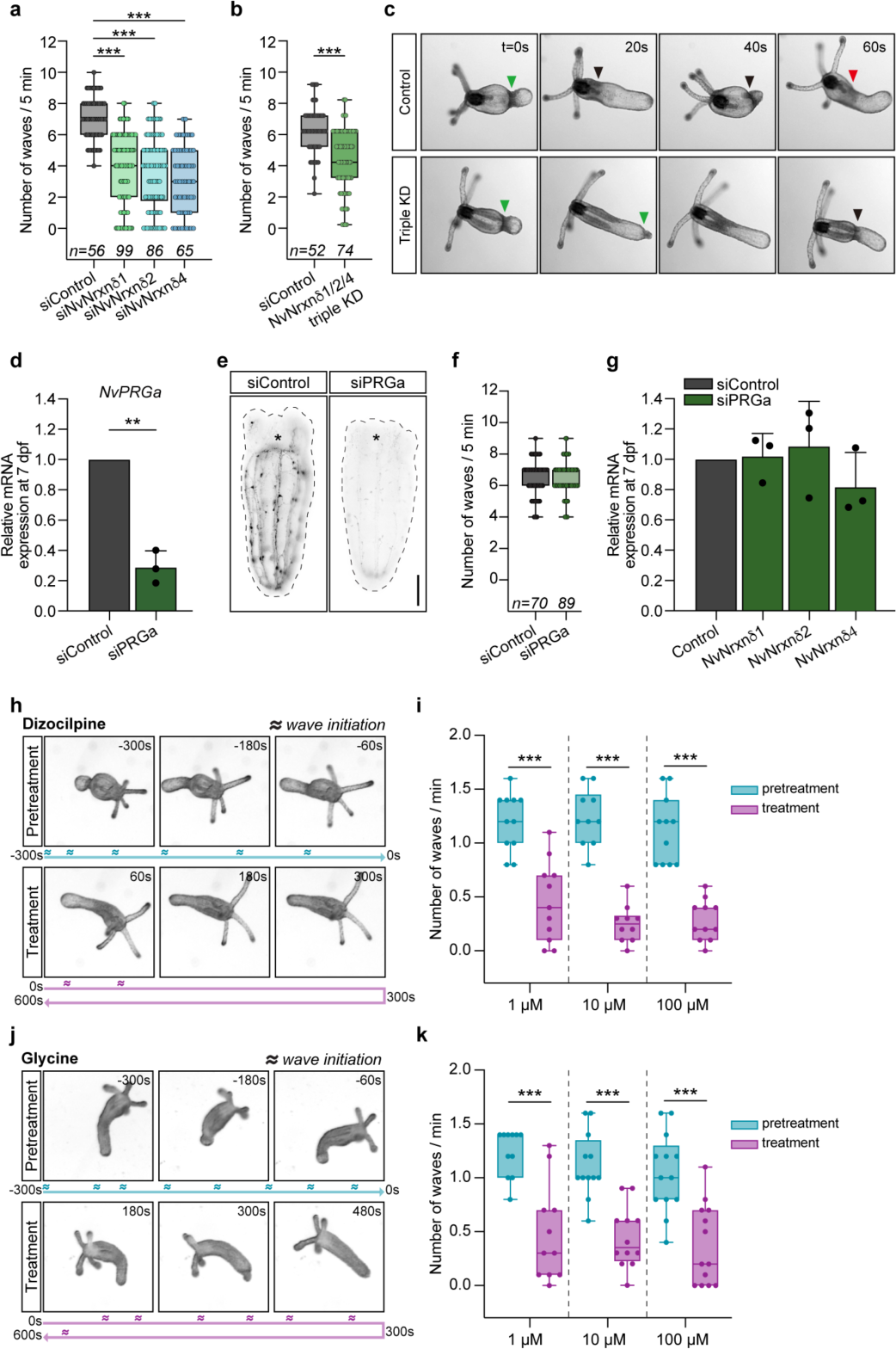
Requirement of δ-Nrxns and glutamate signals for polyp peristalsis. **a** Number of peristaltic waves of polyps transfected with siRNA #2+#3 for NvNrxnδ1, #3 for NvNrxnδ2, or #1+#2 for NvNrxnδ4, respectively. **b** Number of peristaltic waves of polyps after triple knockdown of NvNrxnδ1/2/4 genes by co-transfection of siRNAs (δ1 #3, δ2 #3, and δ4 #2). **c** Representative image time series of peristalsis of control (upper panels) or NvNrxnδ1/2/4 siRNAs-transfected (lower panels) polyps. Green, black, and red arrowheads indicate the 1^st^, 2^nd^ and 3^rd^ propagated wave, respectively. **d** qPCR analysis of NvPRGa mRNA expression of *NvPRGa* siRNA-transfected polyps. **e** Representative immunostaining images of NvPRGamide peptide in *NvPRGa* siRNA-transfected polyps. Scale bar, **100** µm. **f** Peristalsis rate of PRGamide-depleted polyps. **g** Expression of NvNrxnδ genes in PRGamide-depleted polyps. **h-k** Number of peristaltic waves of polyps upon an increasing concentration (1, 10, and 100 µM) of dizocilpine, an ionotropic glutamate receptor NMDA inhibitor (**h, i**), or glycine, an inhibitory neurotransmitter (**j, k**). Two image time series are shown as examples (**h**; 10 µM dizocilpine, **j**; 10 µM glycine). Below the images, wave initiation events are marked as a wave icon on the timeline. mRNA/protein expression and peristalsis were analyzed on 7 dpf and 8 dpf, respectively. **a, b, d, f, g, i, k** Statistical data are presented as means ± SDs for barplots and as interquartiles with minimum and maximum data for boxplots; ** and ***, *p*<0.01 and *p*<0.001, respectively. *P* values were obtained using two-sided unpaired t-tests of at least three independent experiments in **a, b, d, f,** and **q**, or two independent experiments in **i** and **k**.

So how do δ-Nrxns control polyp peristalsis? They do not appear to be expressed at high levels in all PRGamide^+^ neurons (Figure 2c), an indication that they might not participate in PRGamide-mediated peptide neurotransmission. To examine whether peristalsis is modulated by PRGamide neuropeptide-mediated signaling, we decreased PRGamide precursor mRNAs by gene knockdown. Although a significant reduction of PRGamide mRNA levels and immunofluorescent signaling of mature PRGamide peptides was observed upon knockdown (Figure. 3d,e), we did not see a significant effect on the peristaltic wave rate (Figure 3f). Moreover, knockdown of PRGamide did not affect expression of δ-Nrxns (Figure 3g). These data indicate that PRGamide peptidergic signaling is not involved in control of nNrxn-dependent spontaneous peristalsis and support the idea that neuropeptide signals act through synapse-independent volume transmission.

Since δ-Nrxns may not be required in peptidergic signaling, these synaptic cell adhesion molecules may be crucial for signaling mediated by other neurotransmitters. In cnidarians, evidence is accumulating to suggest that glutamate and glycine are responsible for signaling through ionotropic receptors (Pierobon et al. 2001; Kass-Simon et al. 2002; Pierobon et al. 2004). δ-Nrxn^+^ neurons are not only equipped with peptidergic signaling-related genes, but also with molecules necessary for chemical transmission by glutamate and glycine (Hayakawa et al. 2022) (Supplementary Figure 9). We therefore carried out a series of pharmacological assays to determine whether peristalsis is regulated by glutamate and glycine signaling. Interestingly, systemic treatment with L-glutamate or NMDA at concentrations previously used in other cnidarians (Kass-Simon et al. 2003; Pierobon et al. 2001) had no clear effects (Supplementary Figure 10), but the NMDA antagonist, Dizocilpine, resulted in a significant decrease in *N. vectensis* peristalsis (Figure 3h, i). The inhibitory effect on ionotropic GluRs was specific to Dizocilpine, and no significant effect was observed with the Kainate/AMPA receptor antagonist, NBQX (Supplementary Figure 10). Also, treatment with glycine, a major inhibitory neurotransmitter in bilaterians, resulted in a reduction in wave number, similar to inhibition of NMDA-mediated glutamate signaling (Figure 3j, k) and is consistent with the glycine’s inhibitory function observed in other cnidarians (Pierobon et al. 2004; Ruggieri et al. 2004). The above data indicate that spontaneous polyp peristalsis observed in our experimental conditions may be caused by high glutamate signaling activity. Our detection by HCR of genes involved in glutamate and glycine signaling (vesicular transporters: SLC17, SLC32, and SLC6; ionotropic receptors: iGluRs and GlyR alphas) was not successful, probably due to their low expression levels. Therefore, physiological details of the glutamatergic and glycinergic neuronal types in *N. vectensis* remain unclear. Altogether, our results from gene expression patterns, knockdown experiments, and pharmacology suggest that nNrxns regulate peristalsis in *N. vectensis* through classical neurotransmitters such as glutamate, rather than PRGamide peptide signals.

### cNrxn knockdown disrupts proper epithelial organization

After we demonstrated involvement of Nrxn in prebilaterian neuronal function, we decided to examine the ancestral function of Nrxns, which was acquired before the emergence of the nervous system. In cnidarians, cNrxns are a group of broadly-expressed Nrxns that offer insight into preneural functions of Nrxns. In this analysis, we focused on the cNrxn, NvNrxnα3-1, which is mainly expressed in epithelial cells of adult *N. vectensis* (Figure 2a, Supplementary Figure 4). Visualization of mRNA confirmed the epithelial expression of NvNrxnα3-1 during planula and polyp stages (Figure 4a,b). Knockdown of NvNrxnα3-1 caused a significant decrease in mRNA and protein levels without affecting polyp viability (Figure 4c, Supplementary Figure 11). However, knockdown animals displayed decreased epithelial integrity and abnormal morphogenesis (Figure 4d-h). Cortical F-actin staining showed that by the planula stage, ectoderm and endoderm layers were not properly attached to each other (Figure 4d,e, Supplementary Figure 12). On the other hand, staining of Cadherin 1 (NvCdh1), located in the apical and basal junctions of both ectoderm and endoderm layers of *N. vectensis* planula larva (Pukhlyakova et al. 2019), suggested that homophilic cell-cell adhesion in the epithelial layers remains intact (Supplementary Figure 13). The detachment between ectoderm and endoderm caused by NvNrxnα3-1 knockdown became more pronounced at the polyp stage, and an abnormal “blister” appearance of the tentacles was observed (Figure 4f-h). Additionally, some knockdown individuals were unable to achieve normal body size. These findings indicate that cNrxn is involved in proper organization of epithelial tissue through regulation of adhesion between heterogeneous cell types, rather than homologous cell-cell adhesion within each epithelial cell layer.

**Figure 4:**
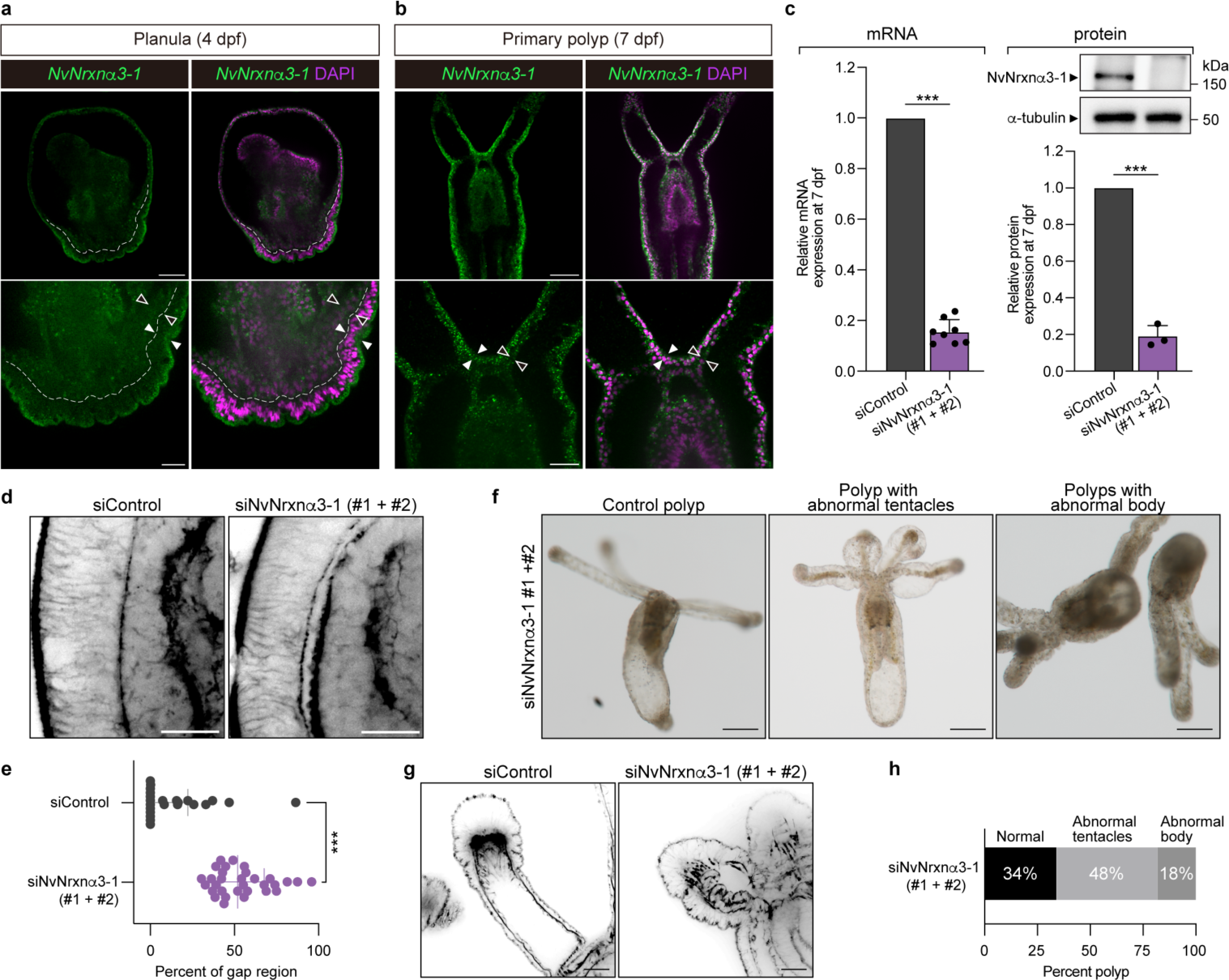
α-Nrxn is required for epithelial organization. **a, b** Representative images of hybridization chain reaction staining of NvNrxnα3-1 mRNA at planula (**a**) and primary polyp (**b**) stages. Lower panels, higher magnification. Dotted line in **a** denotes the ecto/endo boundary. Filled and open arrowheads indicate ectoderm and endoderm, respectively. **c** qPCR and western blotting analysis of 7-dpf polyps transfected with control siRNA or NvNrxnα3-1 siRNAs (#1 + #2). α-tubulin as loading control (right). **d** Representative images pf phalloidin staining (black) of control (left) or NvNrxnα3-1-depleted (right) planulae (4 dpf). **e** Quantification of the epithelial detachment length measured by confocal longitudinal sections of phalloidin-stained planulae (4 dpf). **f** Representative images of typical phenotypes in NvNrxnα3-1depleted polyps: normal, abnormal tentacles, and abnormal body. Polyps with abnormal tentacles have crooked-, or rigid-looking tentacles (middle). Polyps with abnormal bodies have short and underdeveloped body column (right). **g** Representative images phalloidin staining (black) of control (left) or NvNrxnα3-1-depleted (right) polyps. **h** Quantification of the percentages of NvNrxnα3-1 knockdown polyps showing the three phenotypes described in **f**. In **c** statistical data are presented as mean ± SD, and in **e** as median with interquartile range; ** and ***, *p*<0.01 and *p*<0.001, respectively. *P* values were obtained using two-sided unpaired t-test of at least three independent experiments in **c** left, or two independent experiments in **c** right and **e**. Scale bars in **a** and **b** top, **a** and **b** bottom, **d**, **f**, **g**, **50**, **20**, **20**, **100**, **25** µm, respectively.

## DISCUSSION

Previous studies of non-bilaterian nervous systems began with electron microscopic observations of neurons and synapse-like structures. In recent years, the repertoire of neural-related gene homologs in early diverging metazoan genomes and molecular mechanisms of neural development have been analyzed (Putnam et al. 2007; Marlow et al. 2009; Chapman et al. 2010; Nakanishi et al. 2012). Additionally, our understanding of the function of the cnidarian nervous system is rapidly advancing, primarily due to application of molecular biological tools (Ikmi et al. 2014; Servetnick et al. 2017; Cleves et al. 2018; Sanders et al. 2018; Karabulut et al. 2019b; Masuda-Ozawa et al. 2022a). However, despite the high evolutionary conservation of synapse-related molecules in their genomes, there is little direct experimental data on their function in cnidarians; thus, little is clear about pre-bilaterian synaptic functions. Nrxn is a major cell adhesion molecule at presynapses of neurons in bilateral animals and participates in synapse assembly through adhesion with various postsynaptic membrane proteins. Our present study of Nrxns in *N. vectensis* characterizes for the first time in cnidarians the function of a molecule important for synaptic function. We showed that knockdown of neuronal Nrxns resulted in abnormal polyp behavior and demonstrated that Nrxns are involved in regulating motor activity in animals with relatively simple nervous systems.

A genomic survey identified full-length and several lineage-specific short Nrxn genes in early-branching metazoans, further confirming conservation of the presence of putative Nrxn partners (Figure 1c, Supplementary Figures 3-5). In cnidarians, *Heliopora coerulea* (Octocorallia) and *Hydra vulgaris* (Hydrozoa), as well as in early-branching lineages other than cnidarians, sequence conservation of putative Nrxn partners is low, so Nrxn-binding membrane protein candidates could not be predicted. However, the richness of candidate protein repertoires in a wide range of metazoan lineages suggests that their interactions were established before the emergence of neural synapses. New insights were also obtained regarding the ability of Nrxn to form presynaptic complexes via its C terminus. In *N. vectensis* (Anthozoa), short Nrxns (δ- and ε-Nrxns) are the major group of Nrxns bearing a clear C-terminal PDZ motif (Supplementary Figure 1), which is thought to be required for formation of presynaptic complexes in bilaterian neuronal synapses. Our analyses of δ-Nrxn expression and function demonstrated their neural roles in *N. vectensis*, suggesting conservation presynaptic function mediated by the C-terminus. This neurospecific function may also be served by another short Nrxn, the ε type. In all cnidarian species examined, including *N. vectensis*, the ε type also has a PDZ motif (Supplementary Figure 1). Interestingly, ε-Nrxn, the only short Nrxn with a PDZ motif in Hydra, is expressed in their neurons (Supplementary Figure 14). These findings may suggest that acquisition of a PDZ binding motif coincided with the neural functions of nNrxns. In bilaterians, the PDZ binding motif is responsible for forming a complex with the scaffolding proteins like CASK and Mint (Biederer et al. 2000, 2001; Hata et al. 1996). However, CASK and Mint are not exclusively expressed in nNrxn^+^ neurons of *N. vectensis*. This may indicate that the complex was not fully deployed yet in the common ancestor of Cnidaria/Bilateria. Indeed, PDZ motifs are present in cNrxns even in neuronal-free placozoa (Supplementary Fig. 1). Additionally, in *Hydra*, the PDZ motif is conserved in Nrxnα3-1 and 3-2, which are mainly expressed in ectoderm cells (Supplementary Figure 14). Therefore, the presence of the PDZ motif alone is not sufficient to infer the ancestral function of Nrxn at synapses. Attempts to identify the repertoire of Nrxn protein complexes formed in early-branching metazoans, including *N. vectensis*, and the cell types in which they are located, will be important research directions to understand evolutionary processes of synaptic organization and function.

From our results, we could deduce that like the canonical roles of Nrxns in bilaterian neuronal synapses, cnidarian nNrxns are involved in neurotransmission of fast chemical transmitters, e.g., glutamate and glycine, rather than in release of other transmitters, e.g., neuropeptides, which is consistent with previous reports on regulation of peristalsis by acetylcholine signaling (Faltine-Gonzalez et al. 2019). However, much remains to be discovered about mechanisms of action of neurotransmitter signals in cnidarians. Despite the high conservation of nAChR genes, cnidarians appear unable to synthesize acetylcholine (Kass-Simon et al. 2007; Anctil 2009; Chapman et al. 2010; Oren et al. 2014). Although the rich repertoire of GABA(A) receptor genes that are candidate binding partners for Nrxn in *N. vectensis* and their neuronal localization are interesting (Figure 2a), in cnidarians GABA appears to be synthesized primarily in non-neuronal cells (Hayakawa et al. 2022); therefore, its function as a neurotransmitter is still uncertain. Molecular phylogenetic analyzes have also shown that the ability to synthesize many monoamine molecules may have been acquired after bilaterian cladogenesis (Goulty et al. 2023). A clear understanding of the transmitter repertoire and functions in cnidarians may reveal the hidden complexity of their nervous systems, which have been considered simple based on their morphology. Actually, our data provided further insight into the specificity and complexity of cell types or synaptic connections in controlling cnidarian behavior. As seen in the knockdown phenotype, the absence of one or more nNrxns results in the same level of dysregulation. This suggests that the three nNrxn homologs either act cooperatively or have different binding partners and specific pathways to which they belong.

Single-cell transcriptome (SCT) and *in situ* hybridization chain reaction (HCR) also provided new insights into early-branching metazoan cell types that express the most evolutionarily conserved cNrxns. In *N. vectensis* polyps, conserved cNrxns, such as NvNrxnα3-1, are expressed in various cell types, but are most abundant in epithelial cells. Multiple SCT data (Sebe-Pedros, Saudemont, et al. 2018; Steger et al. 2022) have shown neuron clusters that weakly express *N. vectensis* cNrxns. However, in our HCR analysis, we observed undetectable level of cNrxn expression in neurons. This is consistent with experimental data showing that Nrxn protein is abundantly expressed in the epithelia of other sea anemones (Ganot et al. 2015). The fact that cNrxn is expressed in epithelial cells in early-branching metazoans (Supplementary Figures 4,5) suggests that the ancestral function of this cell adhesion apparatus is related to organization of epithelial cell layers. As shown in Figure 4, functional loss of NvNrxnα3-1 in *N. vectensis* caused a wider gap between the ectoderm and endoderm layers. On the other hand, cell adhesion between homogeneous cells in each epithelial cell layer appeared intact. This calls into question the postulated role of cNrxns as a key component of septate junctions (SJs) that function between homogeneous cells. In fact, although cNrxns are highly conserved in early-branching metazoans, the distribution of SJ structures before cnidarians is not clear (Ganot et al. 2015). However, the actual nature of the organization of possibly primordial SJs in these metazoan lineages requires further analysis using definitive SJ marker proteins. Deletion experiments of cNrxns in *N. vectensis* suggested that cNrxns may be involved in adhesion between cells and mesoglia, an extracellular matrix (ECM) formed between the ectoderm and endodermal epithelium. Alternatively, failure of adhesion between both epithelia was observed even in planulae with underdeveloped ECM, indicating that cNrxns may contribute to heterogeneous cell adhesion between ectodermal and endodermal epithelia. In *Hydra*, adhesion between heterogeneous epithelial cells is observed through so-called trans-mesoglea pores (Shimizu et al. 2008). Cell-cell communication between the ectoderm and endodermal epithelium in other cnidarians, including *N. vectensis*, has not yet been reported in detail.

Molecular mechanisms of functions of cNrxns and nNrxns requires further analysis in the future. Figure 5a shows one possible evolutionary scenario for emergence and placement of the Nrxn complex in neurons. If cNrxns serve an important function in cell adhesion between heterologous cells, our data that demonstrate the ancestry and subsequent recruitment of non-neuronal Nrxns into the neural synaptic machinery may offer a hypothesis about the evolution of Nrxn-mediated intercellular contact (Figure 5b). Nrxns were acquired in multicellular ancestral animals and organize adhesion between different cell types, such as ectoderm and endoderm. This new mechanism for regulating cell adhesion between different cell types has diversified in function as the number of cell types increases due to “division of labor” and as multicellular systems become more complex. For example, in the nervous system, division of labor such as sensory and effector cells created a need for specific cell-to-cell communication. Further sophistication of heterogeneous cell adhesion mechanisms in the nervous system through the co-option of Nrxn genes may have been a necessary adaptation toward the evolution from peptide-mediated volume transmission to synaptic transmission mediated by chemical neurotransmitters such as glutamate.

**Figure 5:**
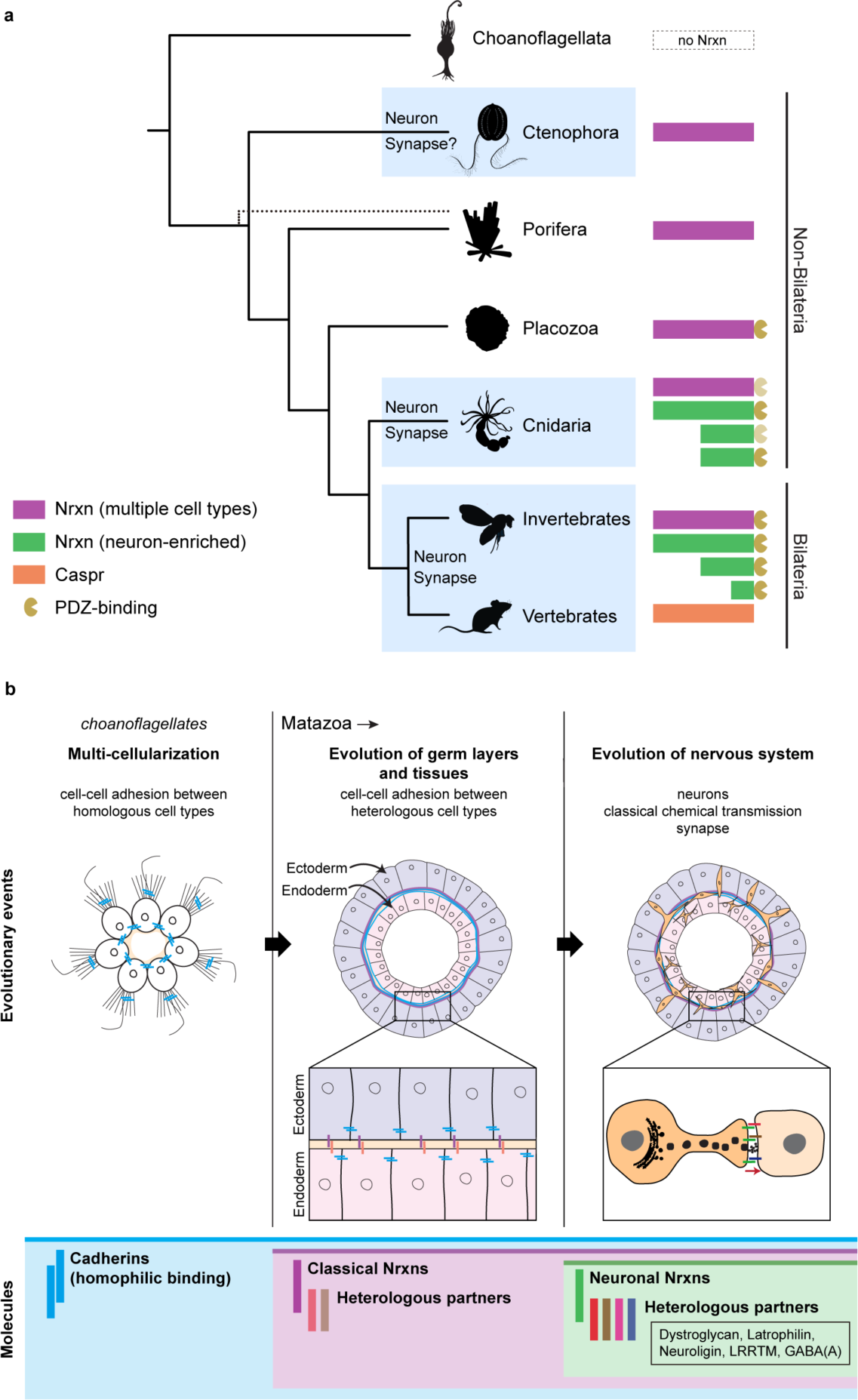
A possible evolutionary scenario of neurexin and neuronal synapses. **a** Nrxns appeared in the last common ancestor of Metazoa. Early Nrxns, which lack the PDZ motif functioned in a variety of cell types, including epithelial cells. Nrxns acquired the PDZ motif in the last common Placozoa/Cnidaria/Bilateria ancestor. Specific neuronal expression and synaptic functions of Nrxn subtypes occurred both in Cnidaria and Bilateria. **b** Proposed hypothesis of the repurposing of Nrxns from heterologous cell-cell adhesion molecules to the chemical synapse organizer. As inferred from the current findings in choanoflagellates, multicellularity in ancestral animals is thought to have begun with colonization of homologous cells. At this evolutionary stage, cells began to be assembled by homophilic binding molecules such as cadherins. Increasing the repertoire of cell types due to division of labor facilitated the diversification of adhesion abilities by employing heterophilic cell adhesion molecules such as Nrxn. In the nervous system, deployment of Nrxns transformed peptide-mediated volume transmission into synaptic transmission, which enables more specific communication with various cell types.

## METHODS

### Protein search

Protein sequences of representative non-bilaterian species from genome or transcriptome studies were obtained (Supplementary Table 1). Synaptic protein sequences were retrieved from UniProtKB/Swiss-Prot database and used as query sequences for Blastp searches. Blastp was run locally using BLAST+ (v2.7.1) or on the NCBI BLAST webpage with an e-value cutoff of 1x10^-5^. Redundant protein sequences that originated from the same gene model were identified and only the longest sequence was included for downstream analysis. Conserved protein domains were identified using consensus predictions of four databases: Conserved Domain Database (CD-Search), ScanProsite (ExPASy), Simple Modular Architecture Research Tool (SMART), and Comprehensive Domain Visualization Tool (CDvist) (Adebali et al. 2015). Conserved motifs and functional sites within a particular domain were checked using ScanProsite. For specific domains, such as PDZ domain, MoDPepInt Server (Kundu et al. 2014) was used to predict PDZ-binding motifs at the C-terminal region of each protein. Transmembrane domains and signal peptides were confirmed with SignalP 4.0 (Petersen et al. 2011) and Phobius (Kall et al. 2007) algorithms. All protein accession IDs retrieved are listed in Supplementary Data 1.

### Protein three-dimensional (3D) modelling

Homology modelling was used to computationally predict the three-dimensional (3D) structures of non-bilaterian Nrxn proteins. Each protein sequence was used as input to the HHPred server (https://toolkit.tuebingen.mpg.de/hhpred) (Soding et al. 2005). Searches were done on the PDB_mmCIF70 database using default settings. The top query-template alignment was converted to PIR file format and forwarded to MODELLER (Sali et al. 1993) for model building. 3D structures were visualized using the UCSF Chimera package (v1.15) (Pettersen et al. 2004). Superimposition was done with the Chimera’s matchmaker command. In addition to MODELLER, the recently available AlphaFold (Jumper et al. 2021) algorithm was also used to further confirm the models.

### Sequence alignments and phylogenetic analyses

Sequence-based analysis of proteins containing repeated domains is particularly difficult. Because of the repeated, shuffled, missing, and highly degenerate domains found in repeat domain proteins, conventional multiple alignment algorithms often fail to yield accurate homology between related proteins. Thus, to avoid possible erroneous output of aligning full-length amino acid sequences containing multiple domain repeats, individual LNS domain sequences were retrieved from each Nrxn homolog. The LNS domains of each protein sequence that aligned against LNS#2 and LNS#6 domains of the template structure were retrieved after the superimposition in Chimera. The LNS#2 and LNS#6 domain sequences from all species were aligned separately using Clustal Omega or the online version of MAFFT-DASH (https://mafft.cbrc.jp/alignment/server/) with default parameters and refinement algorithm L-INS-i (Rozewicki et al. 2019). A python script was written to merge the two alignments. The final alignment containing the LNS#2 and LNS#6 domains (Supplementary Data 2,3) was used to build the phylogenetic tree. Maximum-likelihood analysis was implemented on IQ-TREE (Nguyen et al. 2015) with standard model selection and 5,000 bootstraps. Bayesian inference analysis was executed on the MPI version of MrBayes (v3.2.3) (Ronquist et al. 2012) using two independent Markov chain Monte Carlo (MCMC) runs with four chains per run for 20,000 generations and sample every 100 generations. Phylogenetic trees were visualized in FigTree (v1.4.3) (http://tree.bio.ed.ac.uk/software/figtree/).

### Single-cell expression analysis

Single-cell expression matrices for *N. vectensis* and *M. leidyi* were obtained previously (Hayakawa et al. 2022). The same method was performed to obtain the expression matrices for *T. adhaerens* and *A. queenslandica.* Briefly, protein models were annotated by Blastp alignment to the the *C. elegans* and human UniProt/NCBInr databases with an e-value cutoff of 1x10^-5^. Raw UMI counts, metacell assignments, and metacell annotations were downloaded from http://compgenomics.weizmann.ac.il/tanay/?page_id=724. A global-scaling normalization method was employed to normalize gene expression measurements for each cell by the total expression, multiplied with a scale factor (1,000 for *N. vectensis*; 10,000 for *M. leidyi* and *T. adhaerens*). A single-cell expression plot for *H. vulgaris* was obtained from https://research.nhgri.nih.gov/HydraAEP/ Single Cell Browser.

### Nematostella culture

Animal care, spawning induction, and de-jellying were carried out as previously described (Genikhovich et al. 2009). Briefly, animals were maintained in culture boxes containing 1/3-strength artificial seawater (SEALIFE), hereafter called *Nematostella* medium, and kept in the dark at 18°C. Animals were fed twice per week with freshly hatched *Artemia* (Ocean Star International, Inc.). Spawning was induced by incubating the animals at 26°C for 12-14 hr. Gelatinous mass surrounding the eggs was removed using 4% (wt/vol) L-cysteine (Nacalai Tesque, Inc., cat. 10309-12) in *Nematostella* medium, pH ∼7.5.

### siRNA-mediated knockdown experiments

Custom siRNA design service was provided by Sigma-Aldrich. For each gene, two or three siRNA duplexes were custom synthesized as 21-mers with 3’ dTdT overhangs. To determine the non-specific effects related to siRNA delivery, a non-targeting negative control siRNA with the same modification was also synthesized. All siRNAs used in this study are listed in Supplementary Table 2. siRNA-mediated knockdown was performed as previously described (Karabulut et al. 2019a) with modifications. Briefly, concentrated fertilized eggs were washed three times with fresh *Nematostella* medium. Eggs were transferred to a 2-mL microcentrifuge tube. Then medium was removed and replaced with 6% (wt/vol) Ficoll PM400 (GE Healthcare, cat. GE17-0300-10) in *Nematostella* medium. Each electroporation mixture, containing 200 μL of eggs and siRNAs suspended in 6% Ficoll/*Nematostella* medium, was transferred to a 4-mm-gap sterile cuvette (Bio-Rad, cat. 1652088). Cuvettes were gently tapped for 10 sec on each side to ensure complete mixing of eggs and siRNAs. Eggs were subjected to experimental electroporation parameters (voltage = 50 V; pulse = 25 ms; number of pulses = 1) using the Gene Pulser Xcell Eukaryotic System (Bio-Rad, cat. 1652661). Immediately, cuvettes were filled with *Nematostella* medium and then gently poured into a 150 mm x 15 mm petri dish containing *Nematostella* sperm water. Two control electroporation conditions were included per experiment: 1) with nuclease-free water instead of siRNAs and 2) a non-electroporated condition. Plates were incubated at room temperature (RT).

### RNA extraction, cDNA synthesis, and RT-qPCR

Total RNA was extracted from 5- or 7-day post-fertilization (dpf) stage animals using RNeasy Mini Kit (Qiagen, cat. 74104) according to the manufacturer’s instructions. Genomic DNA residues were removed using an RNase-Free DNase Set (Qiagen, cat. 79254). RNA concentrations were determined using a NanoDrop 2000c spectrophotometer (Thermo Scientific). First strand cDNA synthesis was carried out using the SuperScript IV First-Strand Synthesis System (Thermo Scientific, cat. 18091050). To check siRNA knockdown efficiency, quantitative real-time PCR (qPCR) was conducted on a StepOnePlus Real-Time PCR System (Applied Biosystems) using the PowerUP SYBR Green Master Mix (Applied Biosystems, cat. A25741). All qPCR experiments were performed with a minimum of three technical replicates. Quantification was performed using the –ΔΔCT method with *N. vectensis* GAPDH (XM_001632855.1) as normalization control. Expression levels obtained using GAPDH were comparable with the results using Elongation factor 1-alpha (EF1α) and 18S as control genes. All qPCR primer pairs used in this study are listed in Supplementary Table 3.

### Production of antibodies

For antibodies against NvNrxnα3-1 protein and PRGamide peptides, custom-made polyclonal antibodies were produced in rabbit. The following NvNrxnα3-1 peptides were used for immunization: SFRTYKSSGTVL, LVSQRQMFQYA, MRPPSADLPQN. Each peptide was conjugated with KHL carrier protein. Pooled peptides for NvNrxnα3-1 and the C-terminal structure of amidated peptide (Cys-PRGamide) were used to immunize rabbits with FCA (Freund’s Complete Adjuvant) for the first injection and FIA (Freund’s Incomplete Adjuvant) for boost (Scrum Inc.). After checking immunoreactivity by ELISA assay, anti-sera were affinity-purified using peptide conjugated beads.

### Western blot analysis

Protein extracts were prepared from primary polyps (5 or 7 dpf). Tissues were solubilized in lysis buffer (125 mM Tris-HCl pH 6.8, 2% SDS, 0.01% β-mercaptoethanol, cOmplete Mini EDTA-free (Roche, cat. 04 693 159 001)). After manual homogenization using a plastic pestle, samples were boiled at 95°C for 5 min and sonicated for 15 min. Undissolved samples were separated from the supernatant by centrifugation at 10,000 rpm for 1 min. Protein lysates were stored at - 20°C until western blotting. Prior to SDS-PAGE, protein lysates were prepared by heating at 95°C for 5 min in sample buffer (Nacalai Tesque, Inc, cat. 09499-14). Samples were separated by SDS-PAGE using 7.5% gels or Any kD Mini-PROTEAN TGX Precast Protein Gels (Biorad, cat. 4569033) and transferred to PVDF membranes using a Trans-Blot Turbo Transfer System (Biorad, cat. 1704150). Membranes were incubated with blocking solution (50 mM Tris-HCl pH 6.8, 2 mM CaCl_2_, 80 mM NaCl, 5% skim milk powder, 0.2% NP40) for 1 h at RT. Incubation with primary antibodies (anti-NvNrxnα3-1 -1:1000; anti-alpha-tubulin - 1:1000 (Sigma Aldrich, cat. T6074)) was performed overnight at 4°C. Membranes were washed with wash buffer (PBSTw 0.1%) (3x 10 min each)) before incubation with secondary antibody (anti-rabbit - 1:10,000 (Jackson’s ImmunoResearch, cat. 111-035-003)) for 1 h at RT. After washing (5 x 10 min), detection was performed using the ImmunoStar Zeta (FUJIFILM Wako Pure Chemical Corporation, cat. 291-72401). Band signal quantification was performed using Image Lab software (Biorad).

### Neuropeptide and phalloidin staining

For detection of PRGamide peptides, primary polyps were relaxed using 7% MgCl_2_ in *Nematostella* medium for 10 min and fixed in Zamboni’s fixative (2% paraformaldehyde (PFA), 0.2% picric acid, 0.1M Phosphate buffer, pH 7.2) overnight at 4°C. After fixation, animals were washed in PBS with 0.1% Tween (PBSTw 0.1%) at RT (3x 20 min each), followed by inactivation of residual formaldehyde by a single wash with 0.1M Glycine–HCl pH 7.0 for 15 min at RT. Samples were permeabilized with PBS containing 0.2% Triton X-100 (PBSTr 0.2%) at RT (3x 20 min each). Samples were blocked in blocking buffer (1% BSA, 5% Normal goat serum, 0.1% sodium azide in PBSTw 0.2%) for 1 h at RT. Anti-PRGamide primary antibody was diluted in blocking solution (1:200) and incubation was performed overnight at 4°C, followed by washing in PBSTr 0.2% at RT (3 x 10 min). Animals were then placed in a secondary antibody solution of Alexa Flour 488 AffiniPure Goat Anti-Rabbit IgG (1:500, Jackson ImmunoResearch, cat. 111-545-144) for 1 h at RT. After washing in PBSTr 0.2% at RT (6 x 10 min), drops of SlowFade Gold Antifade Mountant (Invitrogen, cat. S36937) were applied. For F-actin staining with phalloidin, animals were fixed with 4% PFA in PBS for 2 h at 4°C followed by washing with PBSTr 0.2% (4x 20min) at RT. Phalloidin (Cayman Chemical iFluor 488 Conjugate, cat. 20549) and DAPI (Sigma-Aldrich, cat. MBD0015-1ML) were diluted in PBS with 1% BSA and incubated with the samples overnight at 4°C. After washing in PBSTr 0.2% at RT (5 x 15 min), drops of SlowFade Gold Antifade Mountant were applied. Antibodies against *N. vectensis* Cadherin1 (Cdh1) (v1g244010) were graciously shared by Prof. Ulrich Technau (University of Vienna). Planula larvae (3 dpf) were fixed and stained with Cdh1 antibody following (Pukhlyakova et al. 2019) with few modifications. Samples were fixed for 1 h at 4°C with Lavdovsky’s fixative (3.7% formaldehyde (FA), 50% ethanol, 4% acetic acid). After fixation, samples were incubated in ice-cold acetone for 7 min on ice followed by washing with PBSTr 0.2% (5 x 10 min). Samples were then incubated in blocking solution (1% BSA, 5% normal goat serum, 0.1% Sodium azide in PBSTw 0.2%) for 2 h at RT. Anti-Cdh1 antibodies were diluted in blocking solution (1:300) and incubation was performed overnight at 4°C, followed by washing in PBSTr 0.2% at RT (8 x 15 min). After incubation in blocking solution for 2 h at RT, samples were placed in a secondary antibody solution of Alexa Flour 488 AffiniPure Goat Anti-Rat IgG (1:300, Jackson ImmunoResearch, cat. 112-035-003) and DAPI overnight at 4°C. Samples were washed in PBSTr 0.2% at RT (8 x 15 min) and infiltrated with VECTASHIELD PLUS Antifade Mounting Medium (Vector Laboratories, cat. H-1900-2).

### HCR

HCR was performed according to the manufacturer’s instructions (Nepagene, HCR kit, ISHpalette Short hairpin amplifier), with slight modifications. Ten split probes, which were concatenated with initiator sequences for PCR were designed against coding sequences of NvNrxnδ1,-2, and -4, and NvNrxnα3-1 and NvPRGa (Supplementary Data 4). The 4-dpf planulae or 7 dpf polyps of *N. vectensis* were fixed with 4% PFA 4°C overnight and washed with PBSTw 0.1% (3 x 10 min). They were kept in 100% methanol and stored at -20°C until use. These samples were rehydrated with a series of graded MeOH/PBSTw 0.1% washes for 15 min each: 75% MeOH:25% PBSTw 0.1%, 50% MeOH:50% PBSTw 0.1%, 25% MeOH:75% PBSTw 0.1%, and two washes with PBSTw 0.1%. After 30-min treatment with the hybridization solution (Nepagene) for prehybridization treatment, samples were incubated in hybridization buffer containing 20 nM of probes overnight at 37°C. Samples were washed with 0.5x SSC 0.1% Tween20 at 37°C (3 x 15 min), incubated with signal amplification buffer (Nepagene) for 30 min, and with amplification buffer containing hairpin amplifiers (ISH short hairpin amplifier SaraFluor488-S45 and/or ISH short hairpin amplifier ATTO647N-A161: Nepagene) and DAPI for 2 h. Samples were washed with PBSTw 0.1% (3 x 15 min) and mounted on glass slide with SlowFade Gold Antifade Mountant.

### Observation of morphology and behavior

Animal conditions were examined at various developmental stages. For behavior, occurrences of peristaltic waves were counted during the primary polyp stage. A peristaltic wave was defined as a radial contraction of the body column from oral to aboral direction. All waves were counted, including those in progress at the start of the recording and those that did not finish by the end of data collection. Prior to recording, polyps were left to relax their bodies and tentacles. After this, animals were recorded for 5 min and the number of propagated radial contractions during this time was counted. Animals that failed to maintain their orientation, to adhere to the plate, or underwent sudden full-body or tentacle retraction during the 5-min video recording were excluded from these measurements.

### Pharmacology

All experiments were carried out on young polyps (7 to 8 dpf). All reagents were purchased either from Sigma-Aldrich or from Tocris Bioscience. All stock concentrations were prepared by dissolving reagents in Milli-Q water and diluted to their final concentration with *Nematostella* medium. To perform pharmacological assays, polyps were placed in a cell culture plate well (1.9 cm^2^) and left to relax their tentacles. Recordings began 5 min prior to addition of the reagent and continued for an additional 10-min after addition. The number of waves performed by each animal before and after treatment was counted. Only complete, uninterrupted waves were counted, except for those waves at the start and end of recordings. A complete wave is considered when the propagation starts from a position level with pharynx and ends at the foot/physa of the polyp. Control experiments were performed by adding a volume of *Nematostella* medium instead of reagent.

### Imaging

Live images and video recordings were acquired using a stereo microscope (Olympus SZX16). Stained samples were imaged with a fluorescent microscope ECLIPSE Ni (Nikon) equipped with Ds-Ri2 (Nikon) or with SD-OSR microscope system (Olympus). Image and video files were processed with ImageJ/Fiji software.

### F-actin quantification

Calculation of the gap between the ectodermal and endodermal layers was processed using ImageJ/Fiji software as follows. Using phalloidin-stained confocal images of longitudinal sections around the midline of each sample, the length of the basal surface of the ectodermal layer sealed and unsealed from the endodermal epithelium was measured, and the percentage of unsealed length was calculated for each larval sample.

### Statistics and reproducibility

Statistical comparisons were performed using two-tailed unpaired Student’s *t* tests. Data are presented as means ± SDs, scatterplots showing medians with interquartile ranges, or as box plots showing the median (center line), upper and lower quartiles (box limits), and minimum and maximum values (whiskers). *P* values were calculated in GraphPad Prism 9 and Microsoft Excel, and are designated as \**P* < 0.05, \*\**P* < 0.01, and \*\*\**P* < 0.001. Immunostaining and western blot experiments were repeated at least twice, and representative images and blots are shown. All behavior and pharmacological experiments were conducted between 13:00 and 17:00 hr.

## ACKNOWLEDGEMENTS

We are grateful to Akiko Tanimoto and Junko Higuchi for maintaining the *Nematostella* culture, and Ryotaro Nakamura for technical assistance. We thank the OIST Imaging Section for providing access to their microscopes. We also thank Prof. Ulrich Technau (University of Vienna) for the generous gift of anti-NvCdh1 antibodies. This work was supported by OIST Graduate University and by Platform Project for Supporting Drug Discovery and Life Science Research (Basis for Supporting Innovative Drug Discovery and Life Science Research (BINDS)) from AMED under Grant Number JP21am0101110 (support number 0883). HW was supported by JSPS KAKENHI Grant Number JP20K06662. CG was supported by a fellowship from JSPS Research Fellowship for Young Scientists - DC1 (JP19J20655).

## AUTHOR CONTRIBUTIONS

CG and HW conceived and designed the study. CG performed bioinformatic analyses and most of the experiments. CG and KM performed imaging. YK and KT helped with 3D modelling and structure-based sequence alignments. CG, KM, and HW interpreted the results and wrote the manuscript.

## REFERENCES

Adebali, O., D. R. Ortega, and I. B. Zhulin. 2015. ‘CDvist: a webserver for identification and visualization of conserved domains in protein sequences’, Bioinformatics, 31: 1475–7.

Anctil, M. 2009. ‘Chemical transmission in the sea anemone Nematostella vectensis: A genomic perspective’, Comp Biochem Physiol Part D Genomics Proteomics, 4: 268–89.

Biederer, T., and T. C. Sudhof. 2000. ‘Mints as adaptors. Direct binding to neurexins and recruitment of munc18’, J Biol Chem, 275: 39803–6.

Biederer, T., and T. C. Sudhof. 2001. ‘CASK and protein 4.1 support F-actin nucleation on neurexins’, J Biol Chem, 276: 47869–76.

Burkhardt, P., M. Gronborg, K. McDonald, T. Sulur, Q. Wang, and N. King. 2014. ‘Evolutionary insights into premetazoan functions of the neuronal protein homer’, Mol Biol Evol, 31: 2342–55.

Chapman, Jarrod A., Ewen F. Kirkness, Oleg Simakov, Steven E. Hampson, Therese Mitros, Thomas Weinmaier, Thomas Rattei, Prakash G. Balasubramanian, Jon Borman, Dana Busam, Kathryn Disbennett, Cynthia Pfannkoch, Nadezhda Sumin, Granger G. Sutton, Lakshmi Devi Viswanathan, Brian Walenz, David M. Goodstein, Uffe Hellsten, Takeshi Kawashima, Simon E. Prochnik, Nicholas H. Putnam, Shengquiang Shu, Bruce Blumberg, Catherine E. Dana, Lydia Gee, Dennis F. Kibler, Lee Law, Dirk Lindgens, Daniel E. Martinez, Jisong Peng, Philip A. Wigge, Bianca Bertulat, Corina Guder, Yukio Nakamura, Suat Ozbek, Hiroshi Watanabe, Konstantin Khalturin, Georg Hemmrich, André Franke, René Augustin, Sebastian Fraune, Eisuke Hayakawa, Shiho Hayakawa, Mamiko Hirose, Jung Shan Hwang, Kazuho Ikeo, Chiemi Nishimiya-Fujisawa, Atshushi Ogura, Toshio Takahashi, Patrick R. H. Steinmetz, Xiaoming Zhang, Roland Aufschnaiter, Marie-Kristin Eder, Anne-Kathrin Gorny, Willi Salvenmoser, Alysha M. Heimberg, Benjamin M. Wheeler, Kevin J. Peterson, Angelika Böttger, Patrick Tischler, Alexander Wolf, Takashi Gojobori, Karin A. Remington, Robert L. Strausberg, J. Craig Venter, Ulrich Technau, Bert Hobmayer, Thomas C. G. Bosch, Thomas W. Holstein, Toshitaka Fujisawa, Hans R. Bode, Charles N. David, Daniel S. Rokhsar, and Robert E. Steele. 2010. ‘The dynamic genome of Hydra’, Nature, 464: 592–96.

Cleves, Phillip A., Marie E. Strader, Line K. Bay, John R. Pringle, and Mikhail V. Matz. 2018. ‘CRISPR/Cas9-mediated genome editing in a reef-building coral’, Proceedings of the National Academy of Sciences, 115: 5235–40.

Constance, W. D., A. Mukherjee, Y. E. Fisher, S. Pop, E. Blanc, Y. Toyama, and D. W. Williams. 2018. ‘Neurexin and Neuroligin-based adhesion complexes drive axonal arborisation growth independent of synaptic activity’, eLife, 7: e31659.

Craig, A. M., and Y. Kang. 2007. ‘Neurexin-neuroligin signaling in synapse development’, Curr Opin Neurobiol, 17: 43–52.

Dalva, M. B., A. C. McClelland, and M. S. Kayser. 2007. ‘Cell adhesion molecules: signalling functions at the synapse’, Nat Rev Neurosci, 8: 206–20.

Faltine-Gonzalez, D. Z., and M. J. Layden. 2019. ‘Characterization of nAChRs in Nematostella vectensis supports neuronal and non-neuronal roles in the cnidarian-bilaterian common ancestor’, EvoDevo, 10: 27.

Ganot, P., D. Zoccola, E. Tambutte, C. R. Voolstra, M. Aranda, D. Allemand, and S. Tambutte. 2015. ‘Structural molecular components of septate junctions in cnidarians point to the origin of epithelial junctions in eukaryotes’, Mol Biol Evol, 32: 44–62.

Genikhovich, G., and U. Technau. 2009. ‘Induction of spawning in the starlet sea anemone Nematostella vectensis, in vitro fertilization of gametes, and dejellying of zygotes’, Cold Spring Harb Protoc, 2009: pdb prot5281.

Gokce, O., and T. C. Sudhof. 2013. ‘Membrane-tethered monomeric neurexin LNS-domain triggers synapse formation’, J Neurosci, 33: 14617–28.

Gomez, A. M., L. Traunmuller, and P. Scheiffele. 2021. ‘Neurexins: molecular codes for shaping neuronal synapses’, Nat Rev Neurosci, 22: 137–51.

Gorlewicz, A., and L. Kaczmarek. 2018. ‘Pathophysiology of Trans-Synaptic Adhesion Molecules: Implications for Epilepsy’, Frontiers in cell and developmental biology, 6: 119.

Goulty, Matthew, Gaelle Botton-Amiot, Ezio Rosato, Simon G. Sprecher, and Roberto Feuda. 2023. ‘The monoaminergic system is a bilaterian innovation’, Nature Communications, 14: 3284.

Graf, E. R., X. Zhang, S. X. Jin, M. W. Linhoff, and A. M. Craig. 2004. ‘Neurexins induce differentiation of GABA and glutamate postsynaptic specializations via neuroligins’, Cell, 119: 1013–26.

Grayton, Hannah Mary, Markus Missler, David Andrew Collier, and Cathy Fernandes. 2013. ‘Altered Social Behaviours in Neurexin 1α Knockout Mice Resemble Core Symptoms in Neurodevelopmental Disorders’, PLOS ONE, 8: e67114.

Hata, Y., S. Butz, and T. C. Sudhof. 1996. ‘CASK: a novel dlg/PSD95 homolog with an N-terminal calmodulin-dependent protein kinase domain identified by interaction with neurexins’, J Neurosci, 16: 2488–94.

Havrilak, J. A., L. Al-Shaer, N. Baban, N. Akinci, and M. J. Layden. 2021. ‘Characterization of the dynamics and variability of neuronal subtype responses during growth, degrowth, and regeneration of Nematostella vectensis’, BMC Biol, 19: 104.

Hayakawa, E., C. Guzman, O. Horiguchi, C. Kawano, A. Shiraishi, K. Mohri, M. F. Lin, R. Nakamura, R. Nakamura, E. Kawai, S. Komoto, K. Jokura, K. Shiba, S. Shigenobu, H. Satake, K. Inaba, and H. Watanabe. 2022. ‘Mass spectrometry of short peptides reveals common features of metazoan peptidergic neurons’, Nature ecology & evolution, 6: 1438–48.

Ikmi, Aissam, Sean A. McKinney, Kym M. Delventhal, and Matthew C. Gibson. 2014. ‘TALEN and CRISPR/Cas9-mediated genome editing in the early-branching metazoan Nematostella vectensis’, Nature Communications, 5: 5486.

Jekely, G. 2021. ‘The chemical brain hypothesis for the origin of nervous systems’, Philos Trans R Soc Lond B Biol Sci, 376: 20190761.

Jumper, J., R. Evans, A. Pritzel, T. Green, M. Figurnov, O. Ronneberger, K. Tunyasuvunakool, R. Bates, A. Zidek, A. Potapenko, A. Bridgland, C. Meyer, S. A. A. Kohl, A. J. Ballard, A. Cowie, B. Romera-Paredes, S. Nikolov, R. Jain, J. Adler, T. Back, S. Petersen, D. Reiman, E. Clancy, M. Zielinski, M. Steinegger, M. Pacholska, T. Berghammer, S. Bodenstein, D. Silver, O. Vinyals, A. W. Senior, K. Kavukcuoglu, P. Kohli, and D. Hassabis. 2021. ‘Highly accurate protein structure prediction with AlphaFold’, Nature, 596: 583–89.

Kall, L., A. Krogh, and E. L. Sonnhammer. 2007. ‘Advantages of combined transmembrane topology and signal peptide prediction--the Phobius web server’, Nucleic Acids Res, 35: W429–32.

Karabulut, A., S. He, C. Y. Chen, S. A. McKinney, and M. C. Gibson. 2019a. ‘Electroporation of short hairpin RNAs for rapid and efficient gene knockdown in the starlet sea anemone, Nematostella vectensis’, Dev Biol, 448: 7–15.

Karabulut, Ahmet, Shuonan He, Cheng-Yi Chen, Sean A. McKinney, and Matthew C. Gibson. 2019b. ‘Electroporation of short hairpin RNAs for rapid and efficient gene knockdown in the starlet sea anemone, Nematostella vectensis’, Developmental Biology, 448: 7–15.

Kass-Simon, G., A. Pannaccione, and P. Pierobon. 2003. ‘GABA and glutamate receptors are involved in modulating pacemaker activity in hydra’, Comp Biochem Physiol A Mol Integr Physiol, 136: 329–42.

Kass-Simon, G., and P. Pierobon. 2007. ‘Cnidarian chemical neurotransmission, an updated overview’, Comp Biochem Physiol A Mol Integr Physiol, 146: 9–25.

Kass-Simon, G., and Jr A. A. Scappaticci. 2002. ‘The behavioral and developmental physiology of nematocysts’, Canadian Journal of Zoology, 80: 1772–94.

Keijzer, F., and A. Arnellos. 2017. ‘The animal sensorimotor organization: a challenge for the environmental complexity thesis’, Biol Philos, 32: 421–41.

Kundu, K., M. Mann, F. Costa, and R. Backofen. 2014. ‘MoDPepInt: an interactive web server for prediction of modular domain-peptide interactions’, Bioinformatics, 30: 2668–9.

Leshchyns’ka, I., and V. Sytnyk. 2016. ’Synaptic Cell Adhesion Molecules in Alzheimer’s Disease’, Neural Plast, 2016: 6427537.

Li, J., J. Ashley, V. Budnik, and M. A. Bhat. 2007. ‘Crucial role of Drosophila neurexin in proper active zone apposition to postsynaptic densities, synaptic growth, and synaptic transmission’, Neuron, 55: 741–55.

Marlow, H. Q., M. Srivastava, D. Q. Matus, D. Rokhsar, and M. Q. Martindale. 2009. ‘Anatomy and development of the nervous system of Nematostella vectensis, an anthozoan cnidarian’, Dev Neurobiol, 69: 235–54.

Masuda-Ozawa, T., S. Fujita, R. Nakamura, H. Watanabe, E. Kuranaga, and Y. I. Nakajima. 2022a. ‘siRNA-mediated gene knockdown via electroporation in hydrozoan jellyfish embryos’, Sci Rep, 12: 16049.

Masuda-Ozawa, Tokiha, Sosuke Fujita, Ryotaro Nakamura, Hiroshi Watanabe, Erina Kuranaga, and Yu-ichiro Nakajima. 2022b. ‘siRNA-mediated gene knockdown via electroporation in hydrozoan jellyfish embryos’, Scientific Reports, 12: 16049.

Miller, M. T., M. Mileni, D. Comoletti, R. C. Stevens, M. Harel, and P. Taylor. 2011. ‘The crystal structure of the alpha-neurexin-1 extracellular region reveals a hinge point for mediating synaptic adhesion and function’, Structure, 19: 767–78.

Missler, M., W. Zhang, A. Rohlmann, G. Kattenstroth, R. E. Hammer, K. Gottmann, and T. C. Sudhof. 2003. ‘Alpha-neurexins couple Ca2+ channels to synaptic vesicle exocytosis’, Nature, 423: 939–48.

Moroz, L. L., K. M. Kocot, M. R. Citarella, S. Dosung, T. P. Norekian, I. S. Povolotskaya, A. P. Grigorenko, C. Dailey, E. Berezikov, K. M. Buckley, A. Ptitsyn, D. Reshetov, K. Mukherjee, T. P. Moroz, Y. Bobkova, F. Yu, V. V. Kapitonov, J. Jurka, Y. V. Bobkov, J. J. Swore, D. O. Girardo, A. Fodor, F. Gusev, R. Sanford, R. Bruders, E. Kittler, C. E. Mills, J. P. Rast, R. Derelle, V. V. Solovyev, F. A. Kondrashov, B. J. Swalla, J. V. Sweedler, E. I. Rogaev, K. M. Halanych, and A. B. Kohn. 2014. ‘The ctenophore genome and the evolutionary origins of neural systems’, Nature, 510: 109–14.

Moroz, L. L., and A. B. Kohn. 2015. ‘Unbiased View of Synaptic and Neuronal Gene Complement in Ctenophores: Are There Pan-neuronal and Pan-synaptic Genes across Metazoa?’, Integr Comp Biol, 55: 1028–49.

Musser, J. M., K. J. Schippers, M. Nickel, G. Mizzon, A. B. Kohn, C. Pape, P. Ronchi, N. Papadopoulos, A. J. Tarashansky, J. U. Hammel, F. Wolf, C. Liang, A. Hernandez-Plaza, C. P. Cantalapiedra, K. Achim, N. L. Schieber, L. Pan, F. Ruperti, W. R. Francis, S. Vargas, S. Kling, M. Renkert, M. Polikarpov, G. Bourenkov, R. Feuda, I. Gaspar, P. Burkhardt, B. Wang, P. Bork, M. Beck, T. R. Schneider, A. Kreshuk, G. Worheide, J. Huerta-Cepas, Y. Schwab, L. L. Moroz, and D. Arendt. 2021. ‘Profiling cellular diversity in sponges informs animal cell type and nervous system evolution’, Science, 374: 717–23.

Nakanishi, N., E. Renfer, U. Technau, and F. Rentzsch. 2012. ‘Nervous systems of the sea anemone Nematostella vectensis are generated by ectoderm and endoderm and shaped by distinct mechanisms’, Development, 139: 347–57.

Nguyen, L. T., H. A. Schmidt, A. von Haeseler, and B. Q. Minh. 2015. ‘IQ-TREE: a fast and effective stochastic algorithm for estimating maximum-likelihood phylogenies’, Mol Biol Evol, 32: 268–74.

Oren, M., I. Brickner, L. Appelbaum, and O. Levy. 2014. ‘Fast neurotransmission related genes are expressed in non nervous endoderm in the sea anemone Nematostella vectensis’, PLOS ONE, 9: e93832.

Petersen, T. N., S. Brunak, G. von Heijne, and H. Nielsen. 2011. ‘SignalP 4.0: discriminating signal peptides from transmembrane regions’, Nat Methods, 8: 785–6.

Pettersen, E. F., T. D. Goddard, C. C. Huang, G. S. Couch, D. M. Greenblatt, E. C. Meng, and T. E. Ferrin. 2004. ‘UCSF Chimera--a visualization system for exploratory research and analysis’, J Comput Chem, 25: 1605–12.

Pierobon, P., R. Minei, P. Porcu, C. Sogliano, A. Tino, G. Marino, G. Biggio, and A. Concas. 2001. ‘Putative glycine receptors in Hydra: a biochemical and behavioural study’, Eur J Neurosci, 14: 1659–66.

Pierobon, P., A. Tino, R. Minei, and G. Marino. 2004. ‘Different roles of GABA and glycine in the modulation of chemosensory responses in Hydra vulgaris (Cnidaria, Hydrozoa)’, Hydrobiologia, 530: 59–66.

Pukhlyakova, E. A., A. O. Kirillova, Y. A. Kraus, B. Zimmermann, and U. Technau. 2019. ‘A cadherin switch marks germ layer formation in the diploblastic sea anemone Nematostella vectensis’, Development, 146.

Putnam, N. H., M. Srivastava, U. Hellsten, B. Dirks, J. Chapman, A. Salamov, A. Terry, H. Shapiro, E. Lindquist, V. V. Kapitonov, J. Jurka, G. Genikhovich, I. V. Grigoriev, S. M. Lucas, R. E. Steele, J. R. Finnerty, U. Technau, M. Q. Martindale, and D. S. Rokhsar. 2007. ‘Sea anemone genome reveals ancestral eumetazoan gene repertoire and genomic organization’, Science, 317: 86–94.

Ronquist, F., M. Teslenko, P. van der Mark, D. L. Ayres, A. Darling, S. Hohna, B. Larget, L. Liu, M. A. Suchard, and J. P. Huelsenbeck. 2012. ‘MrBayes 3.2: efficient Bayesian phylogenetic inference and model choice across a large model space’, Syst Biol, 61: 539–42.

Rozewicki, J., S. Li, K. M. Amada, D. M. Standley, and K. Katoh. 2019. ‘MAFFT-DASH: integrated protein sequence and structural alignment’, Nucleic Acids Res, 47: W5–W10.

Rudenko, G. 2017. ‘Dynamic Control of Synaptic Adhesion and Organizing Molecules in Synaptic Plasticity’, Neural Plast, 2017: 6526151.

Ruggieri, R. D., P. Pierobon, and G. Kass-Simon. 2004. ‘Pacemaker activity in hydra is modulated by glycine receptor ligands’, Comp Biochem Physiol A Mol Integr Physiol, 138: 193–202.

Ryan, J. F., K. Pang, C. E. Schnitzler, A. D. Nguyen, R. T. Moreland, D. K. Simmons, B. J. Koch, W. R. Francis, P. Havlak, Nisc Comparative Sequencing Program, S. A. Smith, N. H. Putnam, S. H. Haddock, C. W. Dunn, T. G. Wolfsberg, J. C. Mullikin, M. Q. Martindale, and A. D. Baxevanis. 2013. ‘The genome of the ctenophore Mnemiopsis leidyi and its implications for cell type evolution’, Science, 342: 1242592.

Sachkova, M. Y., E. L. Nordmann, J. J. Soto-Angel, Y. Meeda, B. Gorski, B. Naumann, D. Dondorp, M. Chatzigeorgiou, M. Kittelmann, and P. Burkhardt. 2021. ‘Neuropeptide repertoire and 3D anatomy of the ctenophore nervous system’, Current biology: CB, 31: 5274–85 e6.

Sakarya, O., K. A. Armstrong, M. Adamska, M. Adamski, I. F. Wang, B. Tidor, B. M. Degnan, T. H. Oakley, and K. S. Kosik. 2007. ‘A post-synaptic scaffold at the origin of the animal kingdom’, PLOS ONE, 2: e506.

Sali, A., and T. L. Blundell. 1993. ‘Comparative protein modelling by satisfaction of spatial restraints’, J Mol Biol, 234: 779–815.

Sanders, Steven M., Zhiwei Ma, Julia M. Hughes, Brooke M. Riscoe, Gregory A. Gibson, Alan M. Watson, Hakima Flici, Uri Frank, Christine E. Schnitzler, Andreas D. Baxevanis, and Matthew L. Nicotra. 2018. ‘CRISPR/Cas9-mediated gene knockin in the hydroid Hydractinia symbiolongicarpus’, BMC Genomics, 19: 649.

Sebe-Pedros, A., E. Chomsky, K. Pang, D. Lara-Astiaso, F. Gaiti, Z. Mukamel, I. Amit, A. Hejnol, B. M. Degnan, and A. Tanay. 2018. ‘Early metazoan cell type diversity and the evolution of multicellular gene regulation’, Nature ecology & evolution, 2: 1176–88.

Sebe-Pedros, A., B. Saudemont, E. Chomsky, F. Plessier, M. P. Mailhe, J. Renno, Y. Loe-Mie, A. Lifshitz, Z. Mukamel, S. Schmutz, S. Novault, P. R. H. Steinmetz, F. Spitz, A. Tanay, and H. Marlow. 2018. ‘Cnidarian Cell Type Diversity and Regulation Revealed by Whole-Organism Single-Cell RNA-Seq’, Cell, 173: 1520–34 e20.

Senatore, A., T. S. Reese, and C. L. Smith. 2017. ‘Neuropeptidergic integration of behavior in Trichoplax adhaerens, an animal without synapses’, The Journal of experimental biology, 220: 3381–90.

Servetnick, Marc D., Bailey Steinworth, Leslie S. Babonis, David Simmons, Miguel Salinas-Saavedra, and Mark Q. Martindale. 2017. ‘Cas9-mediated excision of Nematostella brachyury disrupts endoderm development, pharynx formation and oral-aboral patterning’, Development, 144: 2951–60.

Shimizu, H., R. Aufschnaiter, L. Li, M. P. Sarras, Jr., D. B. Borza, D. R. Abrahamson, Y. Sado, and X. Zhang. 2008. ‘The extracellular matrix of hydra is a porous sheet and contains type IV collagen’, Zoology (Jena), 111: 410–18.

Soding, J., A. Biegert, and A. N. Lupas. 2005. ‘The HHpred interactive server for protein homology detection and structure prediction’, Nucleic Acids Res, 33: W244–8.

Srivastava, M., E. Begovic, J. Chapman, N. H. Putnam, U. Hellsten, T. Kawashima, A. Kuo, T. Mitros, A. Salamov, M. L. Carpenter, A. Y. Signorovitch, M. A. Moreno, K. Kamm, J. Grimwood, J. Schmutz, H. Shapiro, I. V. Grigoriev, L. W. Buss, B. Schierwater, S. L. Dellaporta, and D. S. Rokhsar. 2008. ‘The Trichoplax genome and the nature of placozoans’, Nature, 454: 955–60.

Srivastava, M., O. Simakov, J. Chapman, B. Fahey, M. E. Gauthier, T. Mitros, G. S. Richards, C. Conaco, M. Dacre, U. Hellsten, C. Larroux, N. H. Putnam, M. Stanke, M. Adamska, A. Darling, S. M. Degnan, T. H. Oakley, D. C. Plachetzki, Y. Zhai, M. Adamski, A. Calcino, S. F. Cummins, D. M. Goodstein, C. Harris, D. J. Jackson, S. P. Leys, S. Shu, B. J. Woodcroft, M. Vervoort, K. S. Kosik, G. Manning, B. M. Degnan, and D. S. Rokhsar. 2010. ‘The Amphimedon queenslandica genome and the evolution of animal complexity’, Nature, 466: 720–6.

Steger, Julia, Alison G. Cole, Andreas Denner, Tatiana Lebedeva, Grigory Genikhovich, Alexander Ries, Robert Reischl, Elisabeth Taudes, Mark Lassnig, and Ulrich Technau. 2022. ‘Single-cell transcriptomics identifies conserved regulators of neuroglandular lineages’, Cell Reports, 40: 111370.

Sudhof, T. C. 2017. ‘Synaptic Neurexin Complexes: A Molecular Code for the Logic of Neural Circuits’, Cell, 171: 745–69.

Sudhof, T. C. 2018. ‘Towards an Understanding of Synapse Formation’, Neuron, 100: 276–93.

Sudhof, T. C. 2021. ‘The cell biology of synapse formation’, The Journal of cell biology, 220: e202103052.

Suga, H., Z. Chen, A. de Mendoza, A. Sebe-Pedros, M. W. Brown, E. Kramer, M. Carr, P. Kerner, M. Vervoort, N. Sanchez-Pons, G. Torruella, R. Derelle, G. Manning, B. F. Lang, C. Russ, B. J. Haas, A. J. Roger, C. Nusbaum, and I. Ruiz-Trillo. 2013. ‘The Capsaspora genome reveals a complex unicellular prehistory of animals’, Nat Commun, 4: 2325.

Takahashi, T. 2020. ‘Comparative Aspects of Structure and Function of Cnidarian Neuropeptides’, Front Endocrinol (Lausanne), 11: 339.

Trotter, J. H., J. Hao, S. Maxeiner, T. Tsetsenis, Z. Liu, X. Zhuang, and T. C. Sudhof. 2019. ‘Synaptic neurexin-1 assembles into dynamically regulated active zone nanoclusters’, The Journal of cell biology, 218: 2677–98.

Uchigashima, M., A. Cheung, J. Suh, M. Watanabe, and K. Futai. 2019. ‘Differential expression of neurexin genes in the mouse brain’, J Comp Neurol, 527: 1940–65.

Zhang, C., D. Atasoy, D. Arac, X. Yang, M. V. Fucillo, A. J. Robison, J. Ko, A. T. Brunger, and T. C. Sudhof. 2010. ‘Neurexins physically and functionally interact with GABA(A) receptors’, Neuron, 66: 403–16.

