## Supplementary figures and tables for "Neuronal and non-neuronal functions of the synaptic cell adhesion molecule, neurexin, in *Nematostella vectensis*"

**Title**

**Evolutionary Neurobiology Unit, Okinawa Institute of Science and Technology Graduate University, Okinawa, Japan**

Christine Guzman, Kurato Mohri, Hiroshi Watanabe

**Artificial Intelligence Research Center, National Institute of Advanced Industrial Science and Technology (AIST), Tokyo, Japan**

Yuko Tsuchiya, Kentaro Tomii

**Present address**

Christine Guzman

Department of Biology, Institute of Zoology, University of Fribourg, CH-1700, Fribourg, Switzerland

*Corresponding author

### SUPPLEMENTARY FIGURES

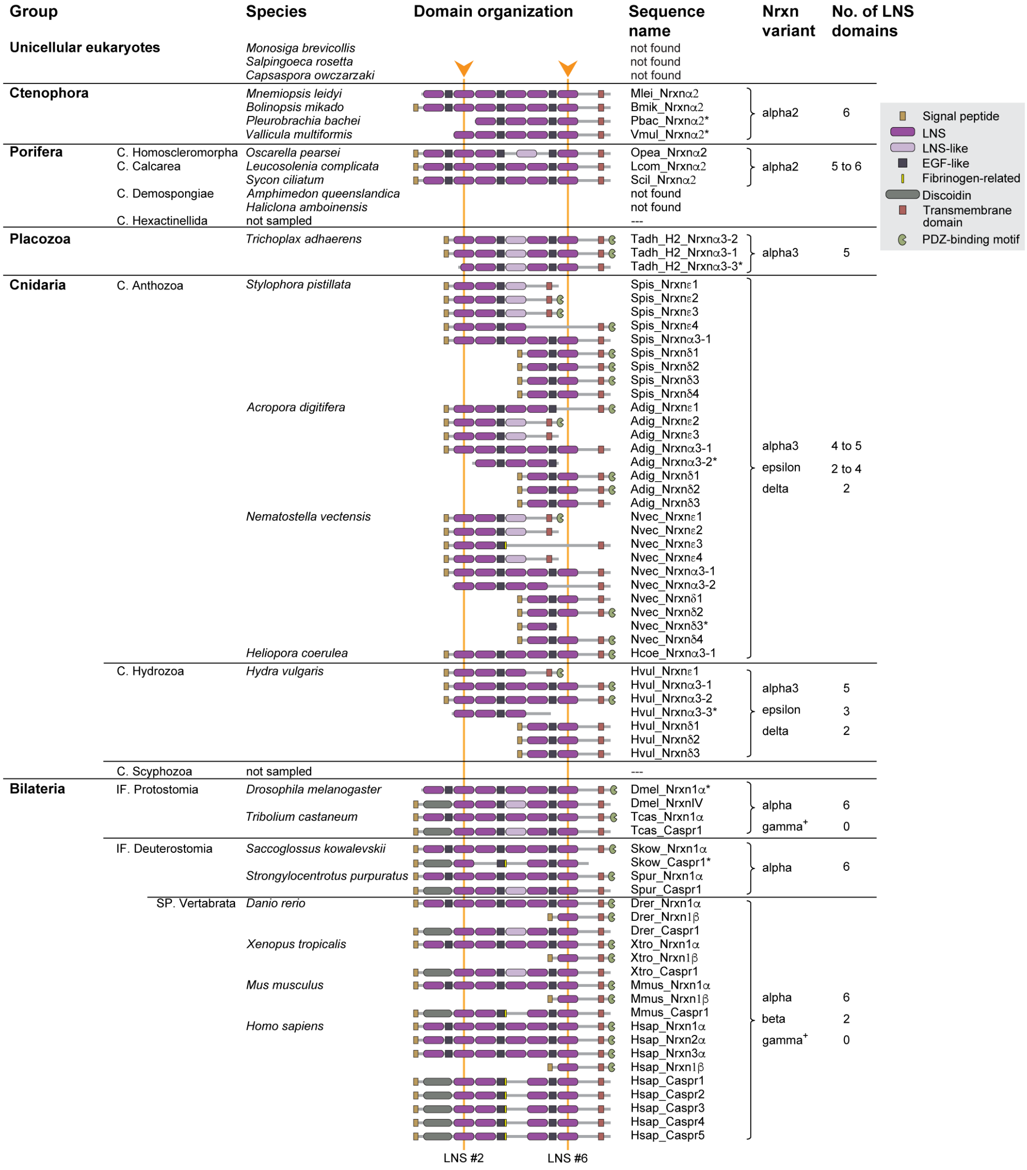


**Supplementary Figure 1.** Protein domain organization of neurexins from different animal groups. The schematic shows which LNS- and EGF-like domains are aligned in the structure analysis. The two most conserved domains (LNS#2 and LNS#6), used for building the multiple sequence alignment in the phylogenetic analysis, are bracketed by orange vertical lines. Incomplete sequences are marked with asterisks. +The schematic for gamma Nrxn is not shown since this newly discovered variant lacks any protein domains. C., Class; IF., Infrakingdom; SP., subphylum.

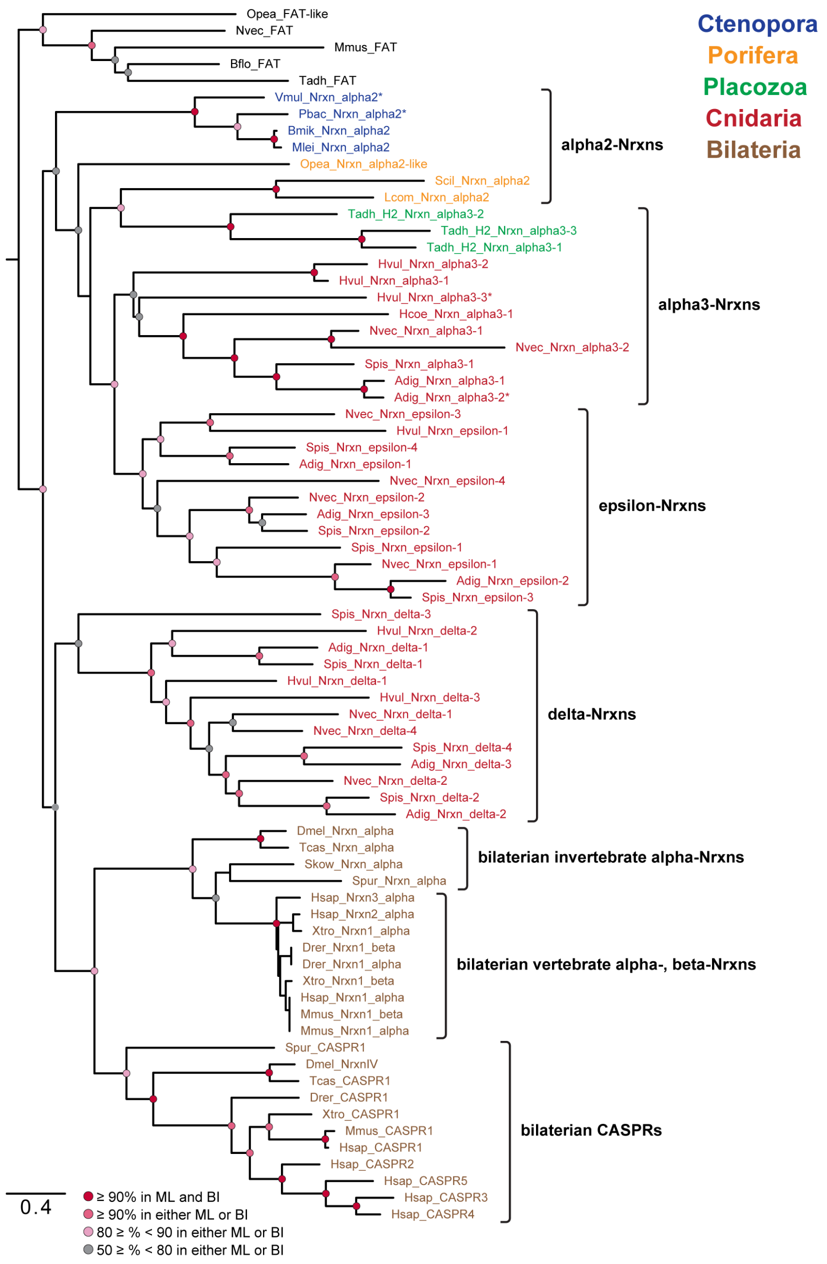


**Supplementary Figure 2.** Phylogenetic tree of the neurexin family. The maximum-likelihood tree was inferred using IQ-TREE with 5,000 bootstraps. A Bayesian tree showed similar branching with the exception of cnidarian epsilon- and delta-Nrxns, in which they are mixed in the same clade. Node support values are derived from bootstrap replicates for the maximum likelihood method, as well as posterior probability for Bayesian analysis. Incomplete sequences are marked with asterisks. The tree was rooted on Fat cadherin proteins. *Nvec, Nematostella vectensis; Adig, Acropora digitifera; Spil, Stylophora pistillata; Hvul, Hydra vulgaris; Hcoe, Heliopora coerulea; Tadh H2, Trichoplax adhaerens H2 strain; Vmul, Vallicula multiformis; Mlei, Mnemiopsis leidyi; Bmik, Bolinopsis mikado; Pbac, Pleurobrachia bachei; Opea, Oscarella pearsei; Scil, Sycon ciliatum; Lcom, Leucosolenia complicata; Aque, Amphimedon queenslandica; Hamb, Haliclona amboinensis; Tcas, Tribolium castaneum; Dmel, Drosophila melanogaster; Skow, Saccoglossus kowalevskii; Xtro, Xenopus tropicalis; Drer, Danio rerio; Mmus, Mus musculus.*

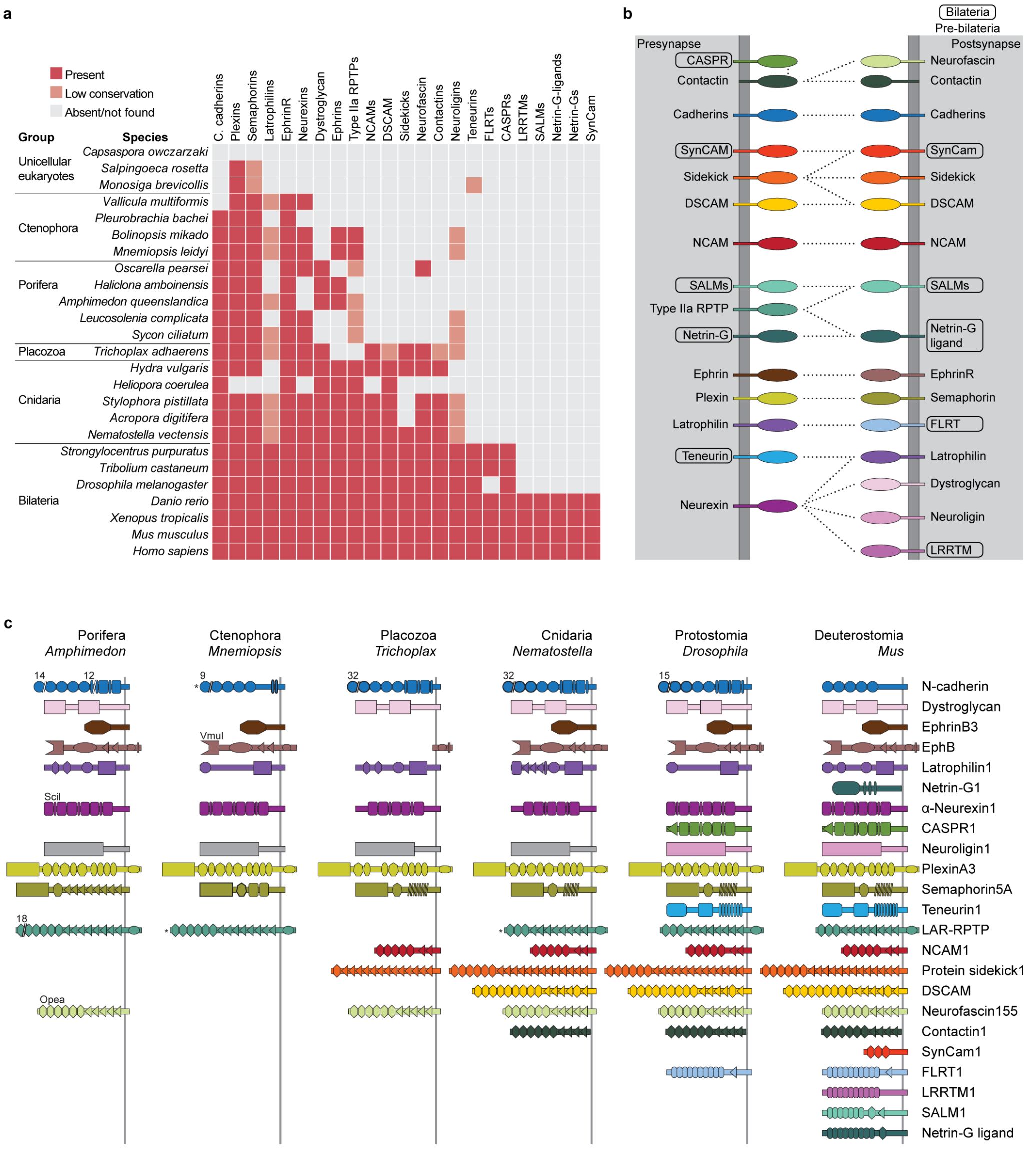


**Supplementary Figure 3.** Phylogenetic distribution of classic synaptic adhesion molecules (SAMs). **a** Presence or absence of SAMs in different metazoan lineages and in unicellular eukaryotes. Complete hits are denoted in red, while partial protein or domain sequences are denoted in orange. Gray boxes indicate absence of the protein. **b** A schematic summary showing whether each SAM originated in non-bilaterians or bilaterians. **c** Schematic protein domain organizations of classic SAMs in different metazoan lineages. The number of repeats is indicated above some protein domains. Asterisks denotes incomplete sequences.

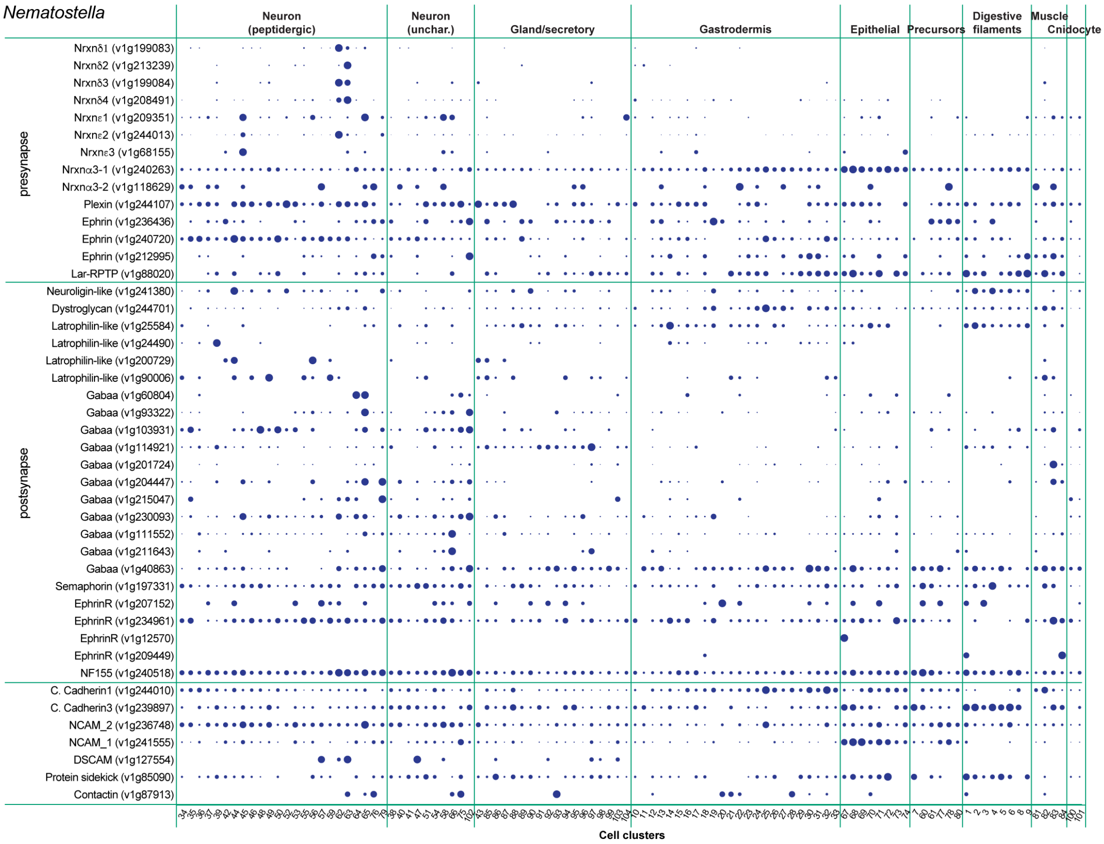


**Supplementary Figure 4.** Abundance of synaptic adhesion molecules (SAMs) and receptors across cell clusters of adult *Nematostella vectensis* (Cnidaria) (Sebe-Pedros et al. 2018a). Dots correspond to normalized UMI count abundance scaled per gene. Dots scale from smallest to largest, corresponding to lowest and highest expression, respectively. Source data are provided as a Source Data file.

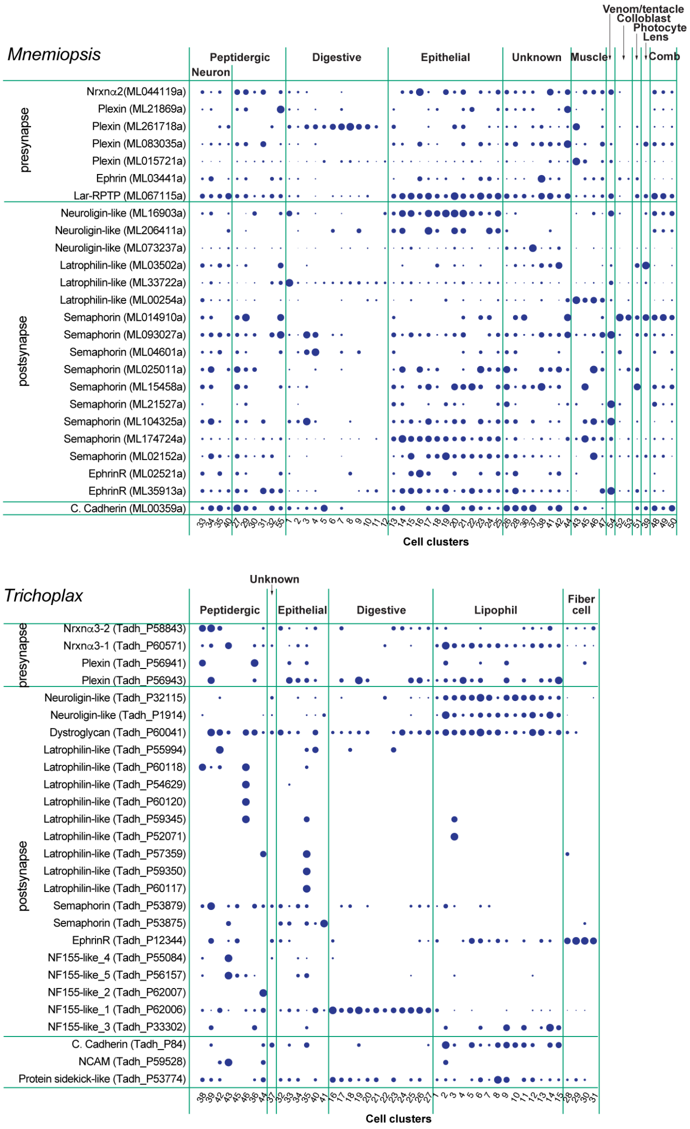


**Supplementary Figure 5.** Abundance of synaptic adhesion molecules (SAMs) and receptors across cell clusters of *Mnemiopsis leidyi* (Ctenophora) and *Trichoplax adhaerens* (Placozoa) (Sebe-Pedros et al. 2018b). Dots correspond to normalized UMI count abundance scaled per gene. Dots scale from smallest to largest, corresponding to lowest and highest expression, respectively. Source data are provided as a Source Data file.

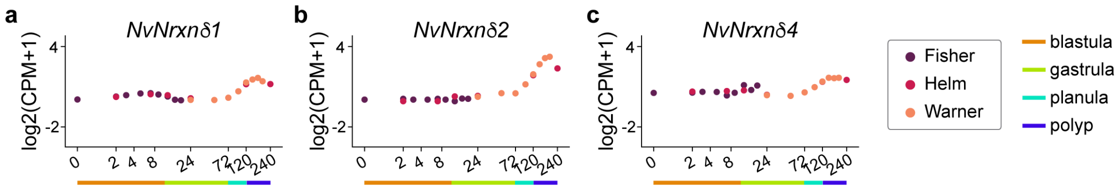


**Supplementary Figure 6.** mRNA expression of *N. vectensis* delta-Nrxns during embryogenesis. All genes begin to be highly expressed at the late planula stage (3 dpf), at which neurons begin to develop neural networks. Data obtained from Warner et al. (2018), Helm et al. (2013), and Fischer et al. (2013). Source data are provided as a Source Data file.

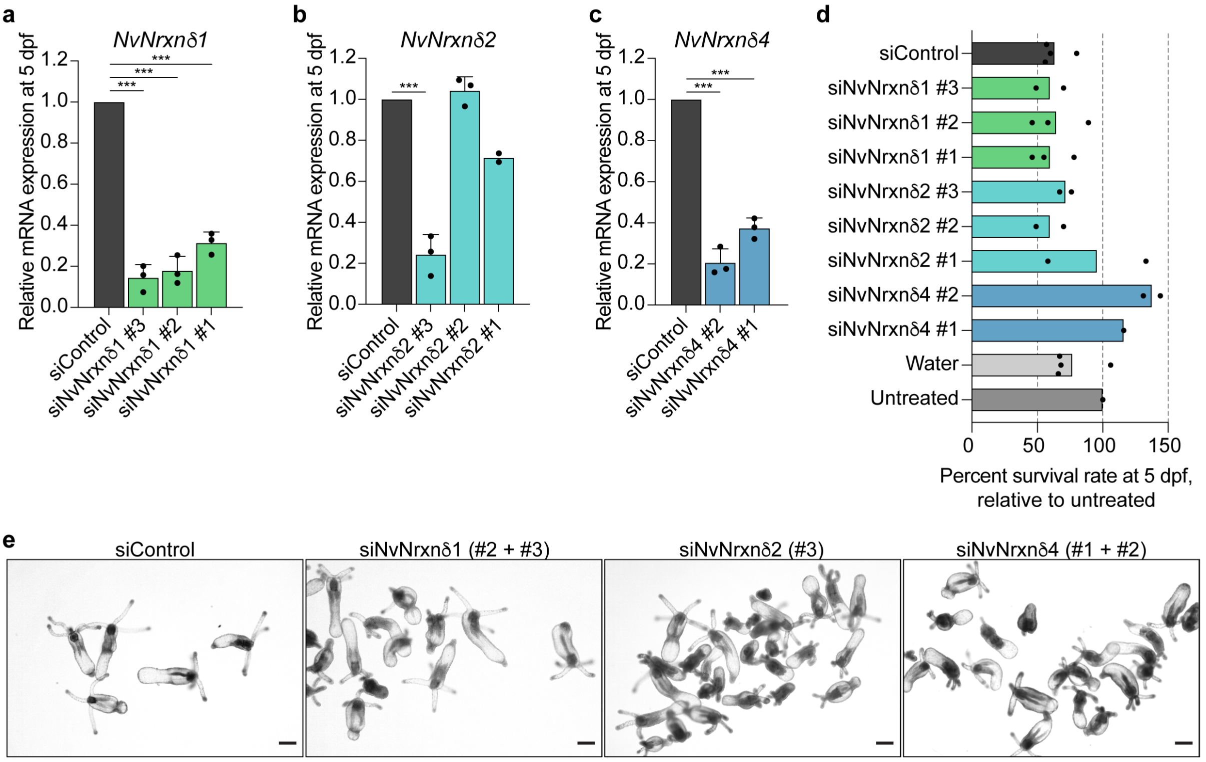


**Supplementary Figure 7. a-c** Knockdown efficiency of siRNAs targeting each delta-Nrxn gene (5 dpf). Expression values are shown relative to samples electroporated with control siRNA. Quantitation was performed using the –ΔΔCT method with *NvGAPDH* as normalization control. *** $p < 0.001$ (Student's t-test). **d** Number of siRNA-electroporated embryos that survived after 5 dpf. **e** No obvious morphological abnormalities were observed in NvNrxnδ1, NvNrxnδ2-, or NvNrxnδ4-knockdown polyps (7 dpf). Scale bars, 100 μm. Source data are provided as a Source Data file.

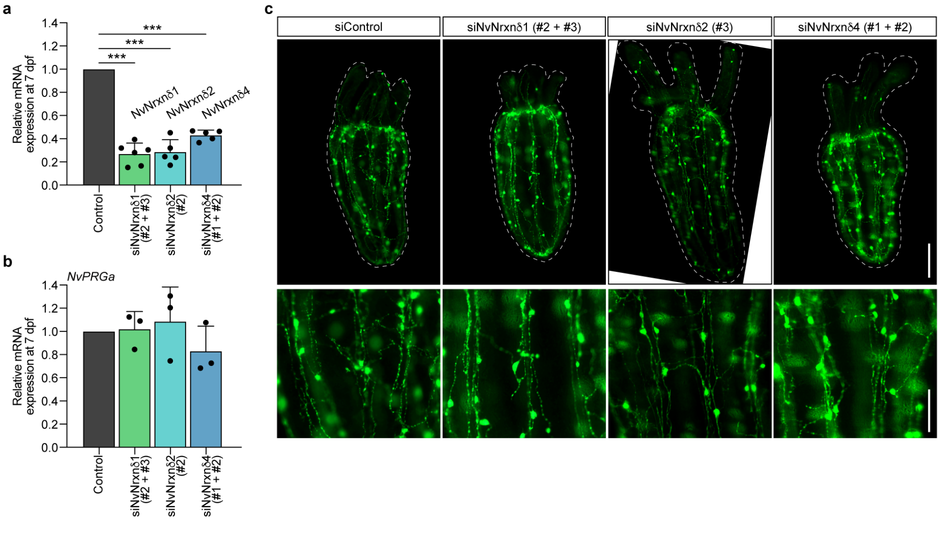


**Supplementary Figure 8.** qPCR analysis of delta-Nrxns (**a**) and NvPRGa (**b**) expression at primary polyp stage (7 dpf) after single gene knockdown (NvNrxnδ1, -2, or -4). Expression values are shown relative to samples electroporated with control siRNA. Quantitation was performed using the –ΔΔCT method with *NvGAPDH* as normalization control. *** $p < 0.001$ (Student's t-test). **c** Representative images of NvPRGamide-positive neurons visualized by immunostaining of 7 dpf polyps transfected with control siRNA or siRNAs for NvNrxnδ1 (#2 + #3), NvNrxnδ2 (#3), and NvNrxnδ4 (#1 + #2). Scale bars, 100 µm. Source data are provided as a Source Data file.

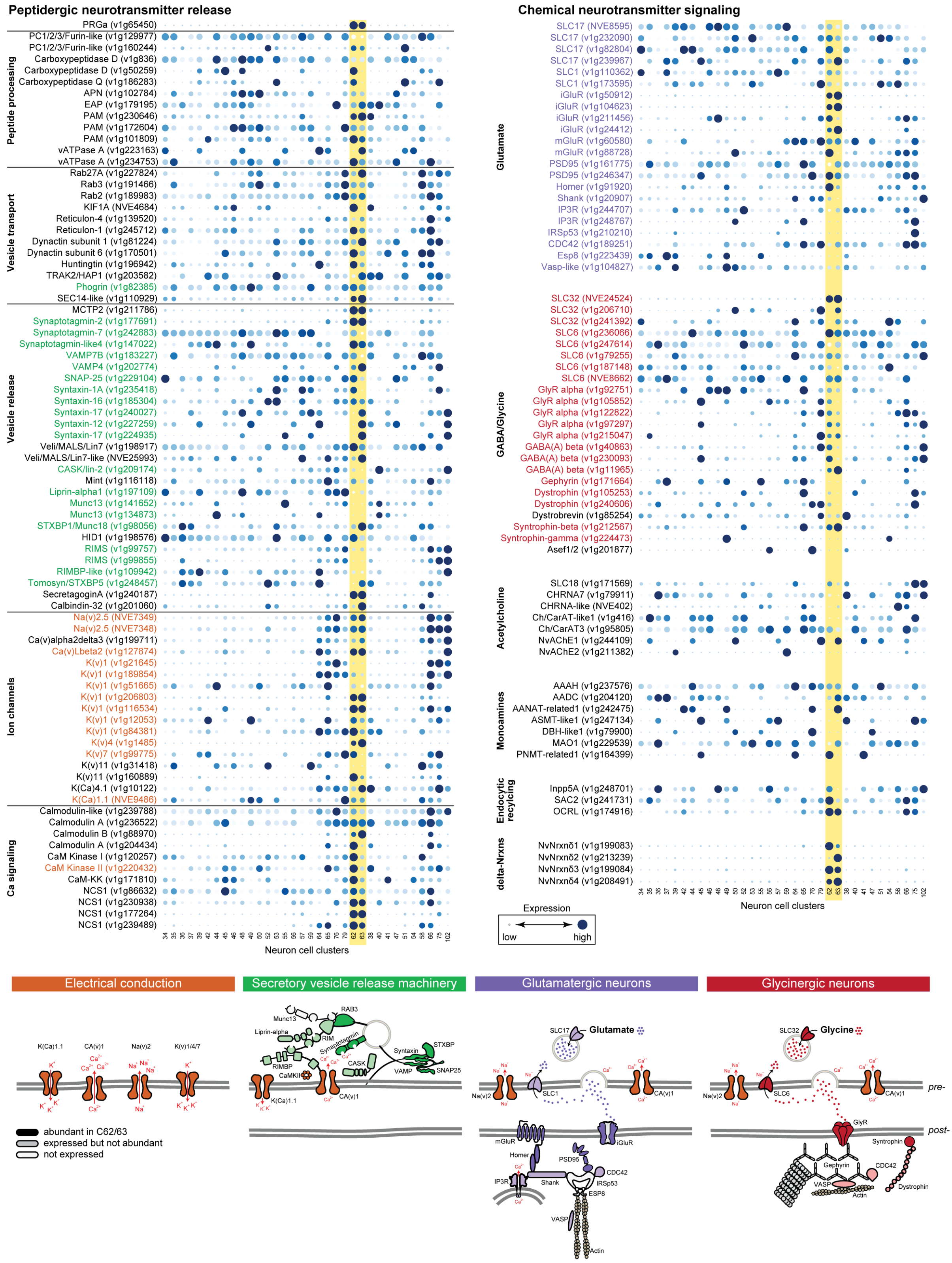


**Supplementary Figure 9.** Abundance of gene homologs involved in neuropeptide and classical neurotransmitter signaling across neuronal cell clusters of adult *N. vectensis* (Sebe-Pedros et al. 2018a). Dots correspond to normalized UMI count abundance scaled per gene. Dots scale from smallest to largest, corresponding to lowest and highest expression, respectively. Schematic representations of the major functional modules required for neuropeptide release and classical neurotransmitter signaling are shown at the bottom panel. Molecules represented in the illustrations are highlighted in dotplots in the upper panel. Molecules exhibiting abundant expression in delta-Nrxn-expressing cell clusters (C62 and C63) are filled in solid colors. Less abundant molecules are semi-transparent. Molecules that are not present in C62 and C63 are empty. Modified schematic representations from Arendt (2020). Source data are provided as a Source Data file.

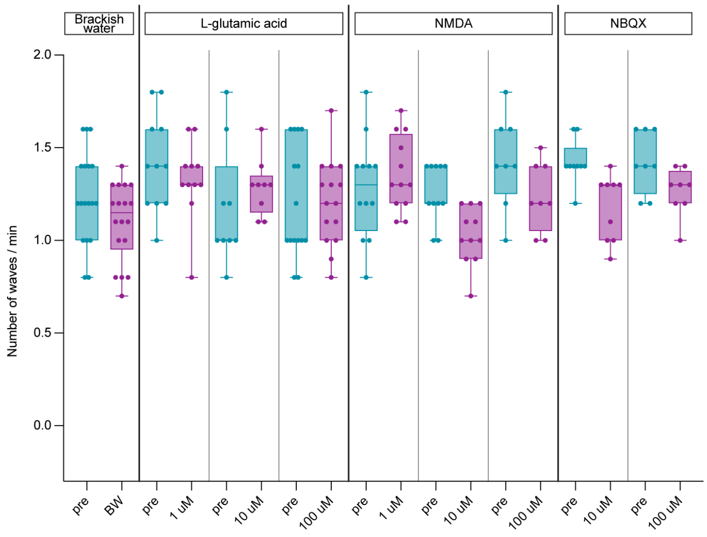


**Supplementary Figure 10.** Number of peristaltic waves of polyps upon increasing concentrations (1, 10, and 100 μM) of L-glutamic acid, NMDA, or NBQX, an AMPA/Kainate receptor inhibitor. Statistical data are presented as interquartiles with minimum and maximum data for boxplots of two independent experiments. Source data are provided as a Source Data file.

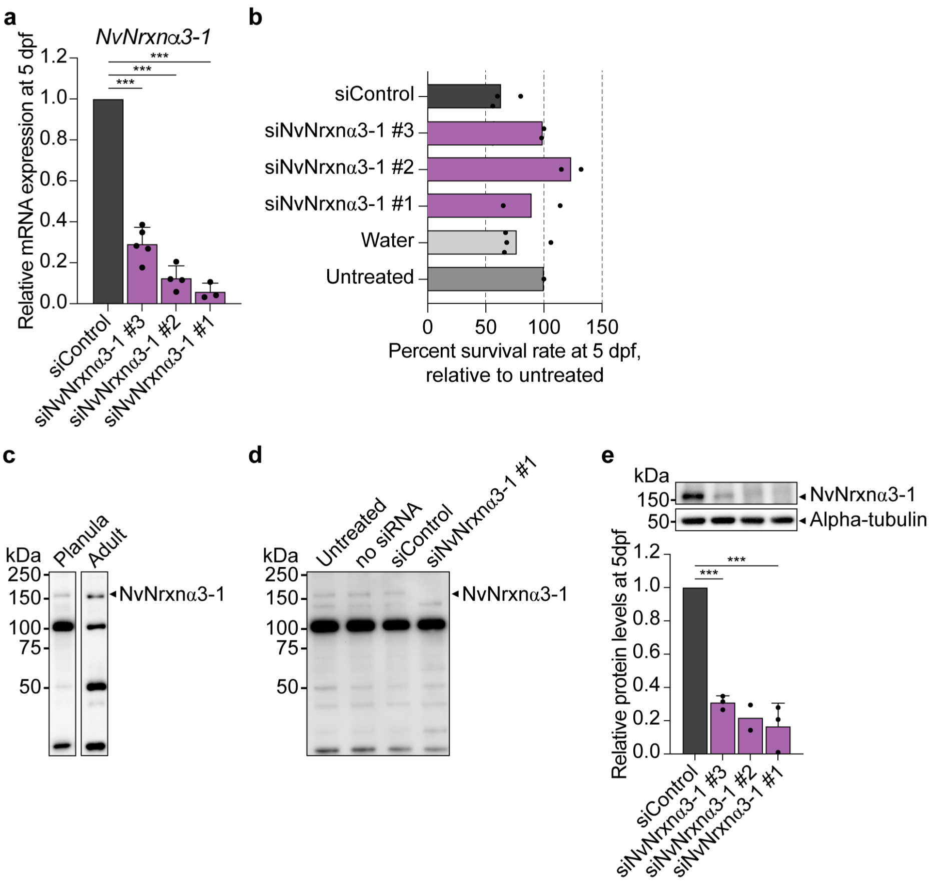


**Supplementary Figure 11. a** Knockdown efficiency of siRNAs targeting NvNrxnα3-1 gene (5 dpf ). Expression values are shown relative to samples electroporated with control siRNA. Quantitation was performed using the –ΔΔCT method with *NvGAPDH* as a normalization control. **b** Relative survival rate of siRNA-electroporated embryos at 5 dpf. **c** Detection of NvNrxnα3-1 protein in planula and adult tissues by western blot analysis. The expected size of NvNrxnα3-1 protein is ~130 kDa, but four distinct bands were observed. **d** Western blot validation at 5 dpf after gene knockdown. The ~150 kDa band was reduced or absent in samples electroporated with siRNA #1 targeting NvNrxnα3-1 mRNA. **e** Protein quantitation using the alpha-tubulin band as a control showed a reduction of protein levels in knockdown animals. *** $p < 0.001$ (Student's t-test). Source data are provided as a Source Data file.

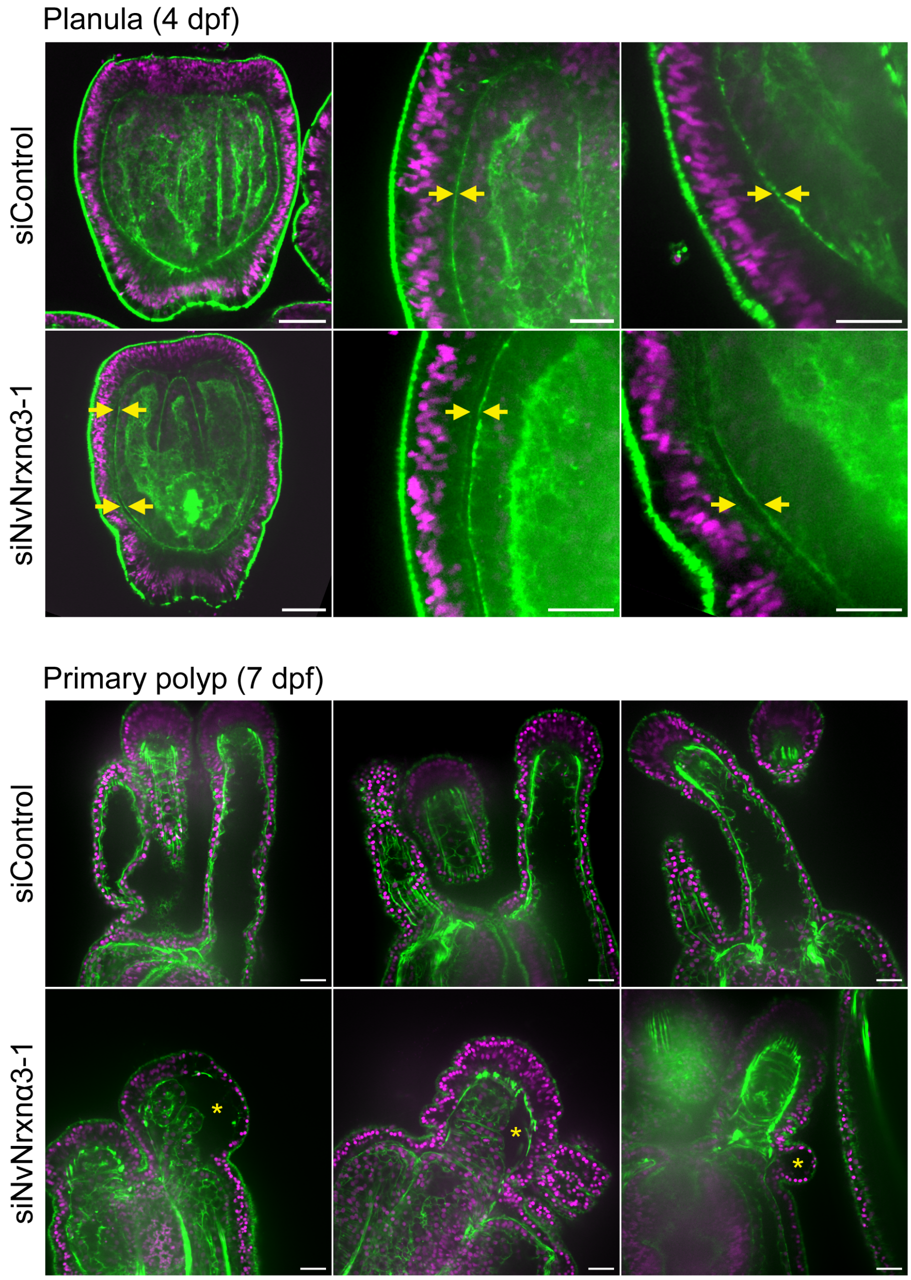


**Supplementary Figure 12.** Representative images of phalloidin staining (green) of control or NvNrxnα3-1-depleted planulae (4 dpf) and primary polyps (7 dpf). Yellow arrows or yellow asterisks in knockdown samples denote a wider gap between ectodermal and endodermal epithelial cells in planulae and primary polyps, respectively. Nuclei are pseudostained in purple (DAPI). Scale bars, 50 μm (top left panels) or 20 μm (others). Source data are provided as a Source Data file.

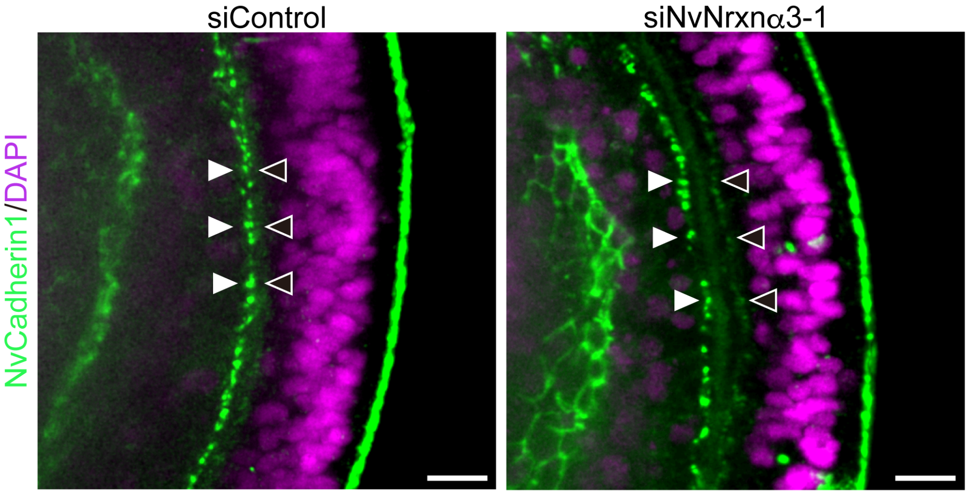


**Supplementary Figure 13.** Cadherin1 staining in control (left) and NvNrxna3-1 knockdown (right) planula larvae. Nuclei are pseudostained in purple (DAPI). Scale bars, 10 μm. Source data are provided as a Source Data file.

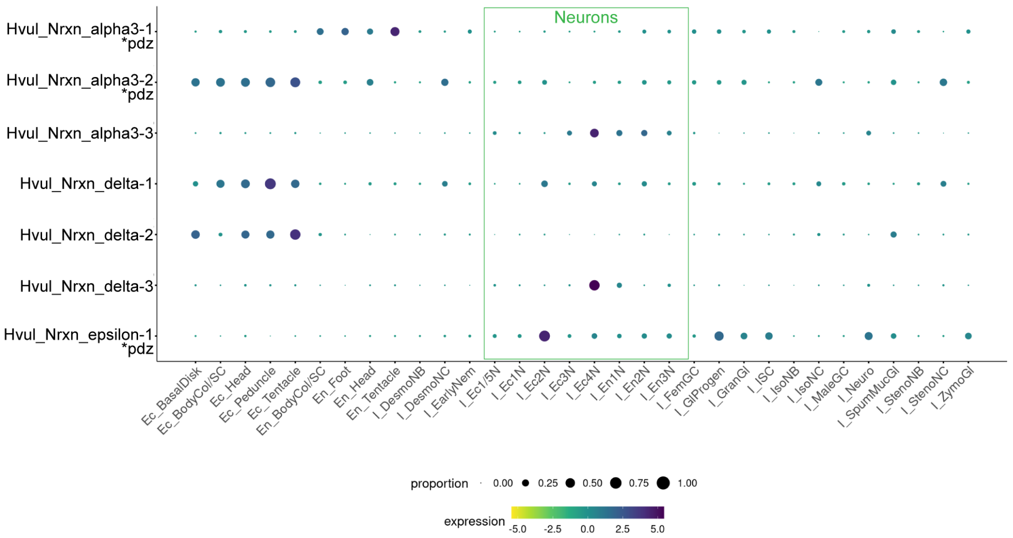


**Supplementary Figure 14.** Single-cell expression plot of Nrxn gene homologs in *Hydra vulgaris* (Cnidaria). Data was obtained from https://research.nhgri.nih.gov/HydraAEP/ Single Cell Browser.

### SUPPLEMENTARY TABLES

**Supplementary Table 1.** List of species used in protein searches.

| Group | Species | Code name | Type of data |
|---|---|---|---|
| Bilateria | *Homo sapiens* | Hsap | Genome |
| | *Mus musculus* | Mmus | Genome |
| | *Xenopus tropicalis* | Xtro | Genome |
| | *Danio rerio* | Drer | Genome |
| | *Strongylocentrotus purpuratus* | Spur | Genome |
| | *Saccoglossus kowalevskii* | Skow | Genome |
| | *Drosophila melanogaster* | Dmel | Genome |
| | *Tribolium castaneum* | Tcas | Genome |
| Cnidaria | *Nematostella vectensis* | Nvec | Genome |
| | *Acropora digitifera* | Adig | Genome |
| | *Stylophora pistillata* | Spis | Genome |
| | *Heliopora coerulea* | Hcoe | Transcriptome |
| | *Hydra vulgaris* | Hvul | Genome |
| Ctenophora | *Bolinopsis mikado* | Bmik | Transcriptome |
| | *Mnemiopsis leidyi* | Mnem | Genome |
| | *Pleurobrachia bachei* | Pbac | Genome |
| | *Vallicula multiformis* | Vmul | Transcriptome |
| Placozoa | *Trichoplax adhaerens, H1* | Tadh | Genome |
| | *Trichoplax adhaerens, H2* | Tadh | Transcriptome |
| Porifera | *Sycon ciliatum* | Scil | Genome |
| | *Leucosolenia complicata* | Lcom | Genome |
| | *Amphimedon queenslandica* | Aque | Genome |
| | *Haliclona amboinensis* | Hamb | Transcriptome |
| | *Oscarella pearsei* | Opea | Genome |
| Unicellular eukaryotes | *Monosiga brevicollis* | Mbre | Genome |
| | *Salpingoeca rosetta* | Sros | Genome |
| | *Capsaspora owczarzaki* | Cowc | Genome |

**Supplementary Table 2.** Target genes and their siRNA designs used in this study.

| Target | RefSeq accession # | Duplex name | siRNA design | |
|---|---|---|---|---|
| | | | Sense strand | Antisense strand |
| *NvNrxnδ1* | XM_001639435.1 | siNvNrxndelta1-1 | GGAAGAACUUCACGGGUAUdTdT | AUACCCGUGAAGUUCUUCCdTdT |
| | | siNvNrxndelta1-2 | GAUGAUCCCGCAUGUGAUUdTdT | AAUCACAUGCGGGAUCAUCdTdT |
| | | siNvNrxndelta1-3 | GGUAUCAUUCAGCAGUUGAdTdT | UCAACUGCUGAAUGAUACCdTdT |
| *NvNrxnδ2* | XM_001628122.1 | siNvNrxndelta2-1 | CUGAUUGGUCCAUGAAAGUdTdT | ACUUUCAUGGACCAAUCAGdTdT |
| | | siNvNrxndelta2-2 | GUUCUAUUAGACAUCAACUdTdT | AGUUGAUGUCUAAUAGAACdTdT |
| | | siNvNrxndelta2-3 | GAAACCACUACUUGUUUAUdTdT | AUAAACAAGUAGUGGUUUCdTdT |
| *NvNrxnδ4* | XM_001631913.1 | siNvNrxndelta4-1 | GAUCUUUCUGGACUCACAAdTdT | UUGUGAGUCCAGAAAGAUCdTdT |
| | | siNvNrxndelta4-2 | CAGCAAAGAUGACCUAGAAdTdT | UUCUAGGUCAUCUUUGCUGdTdT |
| *NvNrxnα3-1* | XM_001637847.1 | siNvNrxnalpha3-1-1 | GAAUCUGUCAUCCGAGAUAdTdT | UAUCUCGGAUGACAGAUUCdTdT |
| | | siNvNrxnalpha3-1-2 | CUGUAACUGUGAUGCUCGUdTdT | ACGAGCAUCACAGUUACAGdTdT |
| | | siNvNrxnalpha3-1-3 | CCAUUCUAUGGCUAUUUCUdTdT | AGAAAUAGCCAUAGAAUGGdTdT |
| *NvPRGa* | XM_001641936.1 | siNvPRG-1 | GAUGAAGAAUCUUUACUUGdTdT | CAAGUAAAGAUUCUUCAUCdTdT |
| Negative control | | siControl | GCAACACGCAGAGUCGUAAdTdT | UUACGACUCUGCGUGUUGCdTdT |

**Supplementary Table 3.** List of qPCR primer pairs used in this study.

| Target | Primer name | Primer sequence |
|---|---|---|
| *NvGAPDH* | NvGapdh_Fw | GGATGGACCAAGTGCCAAGAAC |
| | NvGapdh_Rv | GCTTGCCGTTTACCTCAGGAATGA |
| *NvNrxnδ1* | Nv_deltaNrxn1_For | CCCTCACAGTTATTTTCTCCTC |
| | Nv_deltaNrxn1_Rev | GGTTCGAAGGTCGTATATGAGT |
| *NvNrxnδ2* | Nvec_deltaNrxn2_For | TCTTATGTGGTCTGGTACTTGC |
| | Nvec_deltaNrxn2_Rev | TGGAGCTTCGTACTGAGAGTAA |
| *NvNrxnδ4* | Nvec_deltaNrxn4_F | GCTGTACCTAAACCAGAGCAGT |
| | Nvec_deltaNrxn4_R | GACCGAGTTGGGATAGTTGTAT |
| *NvNrxnα3-1* | Nv_alphaNrxn3-1_F | GATGTCAAGGAGACGTGTGA |
| | Nv_alphaNrxn3-1_R | GACTTGTATGTGCGGAATGA |
| *NvPRG* | PRGamide_Fw | GCAGGTCCCTTATTGAGCTTC |
| | PRGamide_Rv | GTCCGACTTCTCAGCAGACC |
| *Nv18S* | Nv18S-1 Fw | TGGATGACCTCTTTGGCACC |
| | Nv18S-1 Rv | GTGCCCTTCCGTCAATTCCT |
| *NvEF1a* | NvEf1alpha_Fw | GGTTGCCTCTTCGCTTACCACT |
| | NvEf1alpha_Rv | CGTTCCTGGCTTTAGGACAC |

**References**

Sebe-Pedros, A., et al., Cnidarian Cell Type Diversity and Regulation Revealed by Whole-Organism Single-Cell RNA-Seq. Cell, 2018a. 173(6): p. 1520-1534 e20.

Sebe-Pedros, A., et al., Early metazoan cell type diversity and the evolution of multicellular gene regulation. Nat Ecol Evol, 2018b. 2(7): p. 1176-1188.

Arendt, D., The Evolutionary Assembly of Neuronal Machinery. Curr Biol, 2020. 30(10): p. R603-R616.

Warner, J.F., et al., NvERTx: a gene expression database to compare embryogenesis and regeneration in the sea anemone Nematostella vectensis. Development, 2018. 145(10).

Helm, R.R., et al., Characterization of differential transcript abundance through time during Nematostella vectensisdevelopment. BMC genomics, 2013. 14(1): p. 1- 10.

Fischer, A.H. and Smith, J., Nematostella High-density RNAseq time-course. 2013.
